# Linear scaling reveals low-dimensional structure in observable microbial dynamics

**DOI:** 10.1101/2025.06.13.659614

**Authors:** Zhengqing Zhou, Xiaoli Chen, Emrah Şimşek, Grayson S. Hamrick, Yasa Baig, Zachary A. Holmes, Zhenjiao Du, David K. Karig, Lingchong You

## Abstract

Microbial communities often exhibit apparently complex dynamics driven by myriad interactions among community members and with their environments. Yet, practical modeling and control are often based on limited number of observables, raising a fundamental question: *to what extent are these observed dynamics predictable given unobserved background complexity?* Here, we report an emergent simplicity that the temporal dynamics of observable microbial populations can be captured by low- dimensional representations. Using variational autoencoders (VAEs), we define a critical latent dimension (*E****c***) that quantifies the minimal number of variables required to represent observable microbial dynamics. We find that *Ec* scales linearly with the number of observables, despite the complexity of unobserved background dynamics. This principle holds across simulations of ecological, spatial, and gene-transfer models, experiments with engineered and environment-derived communities, and human microbiomes. Our findings establish a scaling law for microbial community dynamics and demonstrate observable dynamics alone contain sufficient information for prediction and control, even without full knowledge of the community.

## Introduction

Microbial communities can harbor hundreds to thousands of species^1^ that engage in interactions on different levels. At the metabolic level, microbes compete for resources and undergo cross-feeding by sharing metabolites^2^. Genetically, they spread functional genes through horizontal transfer^3^. At the population level, they engage in chemical-based communication that controls collective functions, such as virulence^4^ and biofilm formation^5^. These interactions often unfold in structured environments^6–8^, undergo environmental fluctuations^9–11^, and are further shaped by feedbacks from microbial modulation of their surroundings^12,13^.

As a result, both natural and synthetic communities can exhibit non-trivial dynamics like abrupt community shifts^14^ and oscillations^15,16^. The sheer number of interacting components, combined with environmental fluctuations, creates an apparent curse of dimensionality: in principle, fully predicting or controlling the system would require tracking every strain, metabolite, gene flow event, and interaction across time and space. In practice, this level of resolution is experimentally prohibitive to generate and computationally daunting to analyze, as the number of model parameters required to describe such systems grows combinatorially. To manage this complexity, researchers often resort to application of simplification techniques, including dimension reduction^10,14,17^, grouping of sequence variants^18^ and metabolites^19,20^, removal of low-abundance taxa^18,21,22^, and coarse-graining interactions into statistical co-occurrence patterns^23^ or pairwise interaction coefficients in a general Lotka-Volterra model (gLV) ^15,21,22,24–26^. While these simplifications are necessary for specific model formulation and data analysis, it is unclear whether and to what extent the methods or the underlying assumptions are readily generalizable to other microbial ecosystems.

Within this high-dimensional complexity, however, researchers often focus on a small subset of focal populations or functions that are relevant for ecological understanding or biomedical applications. Examples include invasive pathogens that disrupt host health^27–29^, plasmids encoding antibiotic resistance^30^, engineered probiotics as therapeutics^31,32^ and biosensors^33,34^, short-chain fatty acid production in engineered microbiome^35^, and commensal health-associated strains^36^. Under such scenarios, researchers often track the temporal dynamics of focal populations with label-based methods^27–29,31–33,37^. In contrast, the profiling of the entire community based on 16S rRNA sequencing can be costly, and typically provides only genus-level resolution, obscuring strain-level or functional dynamics. As a result, focusing on a small set of observables becomes both practically necessary and biologically meaningful in complex communities.

However, these observables are not isolated; they are dynamically entangled with the broader, unmeasured background community. This entanglement raises a fundamental question: *what information is essential to enable the prediction and control of the observed dynamics, given the unobserved background complexity?*

Evidence suggests that microbial dynamics may reside in a low-dimensional space, governed by a small number of variables. For instance, without explicitly modelling the gut microbiota composition or detailed immune features, a 7-variable ordinary differential equation (ODE) model was sufficient to reproduce the co-infection dynamics of wildtype and mutant *Yersinia enterocolitica* across different mouse hosts^38^. A gLV model using the 18 most prevalent genera captured the temporal patterns during preterm infant gut microbiota assembly, and identified key interaction motifs within the communities^22^.

Dynamic coexistence patterns were captured by taking growth rate as a function of biomass, assuming the community dynamics can be described by a single ecological coordinate^39^. Empirical dynamic modelling (EDM) has been used to forecast the microbial dynamics by constructing low-dimensional, time-delay embeddings of single amplicon sequence variant (ASV) time series^14^. The temporal changes in the microbiome under dietary and antibiotic perturbations could be decomposed into several common temporal patterns^40^. These results suggest the observable microbial dynamics can be represented by a few key variables, enabling predictions despite the complexity of the full system. Despite empirical evidence, a systematic quantification of the low dimensionality of the observable microbes remains absent.

We developed a data-driven approach by using variational autoencoders (VAEs) to extract effective representation and examine the essential dimensionality of observed microbial community dynamics. VAEs are neural network models that can compress complex data, including images^41^, single cell RNA sequence data^42^, spatial profiles^43^, and time-varying data^44,45^ into a latent space, enabling dimensionality reduction and high-fidelity reconstruction. Our previous studies have demonstrated the use of autoencoders to facilitate small chemical detection^45^, simulation acceleration^43^, microbial phenotype classification and parameter estimation^44^. Building on this foundation, we use the VAE as a quantitative tool to probe the minimum number of latent variables, what we term the “critical dimension” (*Ec*), needed to achieve compact and high-fidelity representation of the observed microbial dynamics. Our approach is purely data-driven, predictive, and flexible with complex temporal patterns even subject to environmental perturbations.

Using this framework, we analyzed microbial community dynamics across a broad range of simulated and experimental systems. From numerical simulations of different microbial ecosystems, we quantified the relationship between the number of observables and their critical dimension. Across these ecosystems, we found that *Ec* increases linearly with the number of observables, regardless of the complexity of unmeasured background species, interaction networks, or ecological processes. We experimentally validated this scaling law in plasmid-encoded fluorescence dynamics using synthetic communities consisting of 62 barcoded *E. coli* Keio strains. The scaling law holds for 5 experimental communities from environment-derived inoculum, and longitudinally tracked human vaginal microbiome. We further demonstrate the use of the low-dimensional embedding for forecasting the target microbe abundance and community compositional shifts.

## Results

### VAE reveals low-dimensional structure of observed microbial dynamics in simulations

To determine the effective representation of the observables, we first generated simulated microbial community dynamics using gLV models^8,15,26^ (**Methods**). In each simulation of *N* (=100)-member randomized microbial community, we selected the first *n* members as observables and treated the remaining *N*-*n* members as unobserved background populations (**Fig. 1a**). In each simulation, we randomized the initial abundances of the *n* observables while fixing the initial abundances of the background populations across simulations, but away from equilibrium. We refer to this simulation configuration as the “fixed” case. Since the observables interact with the non-equilibrium backgrounds, their dynamics drastically differed from simulations when they were isolated (**Fig. S1**).

**Figure 1.**
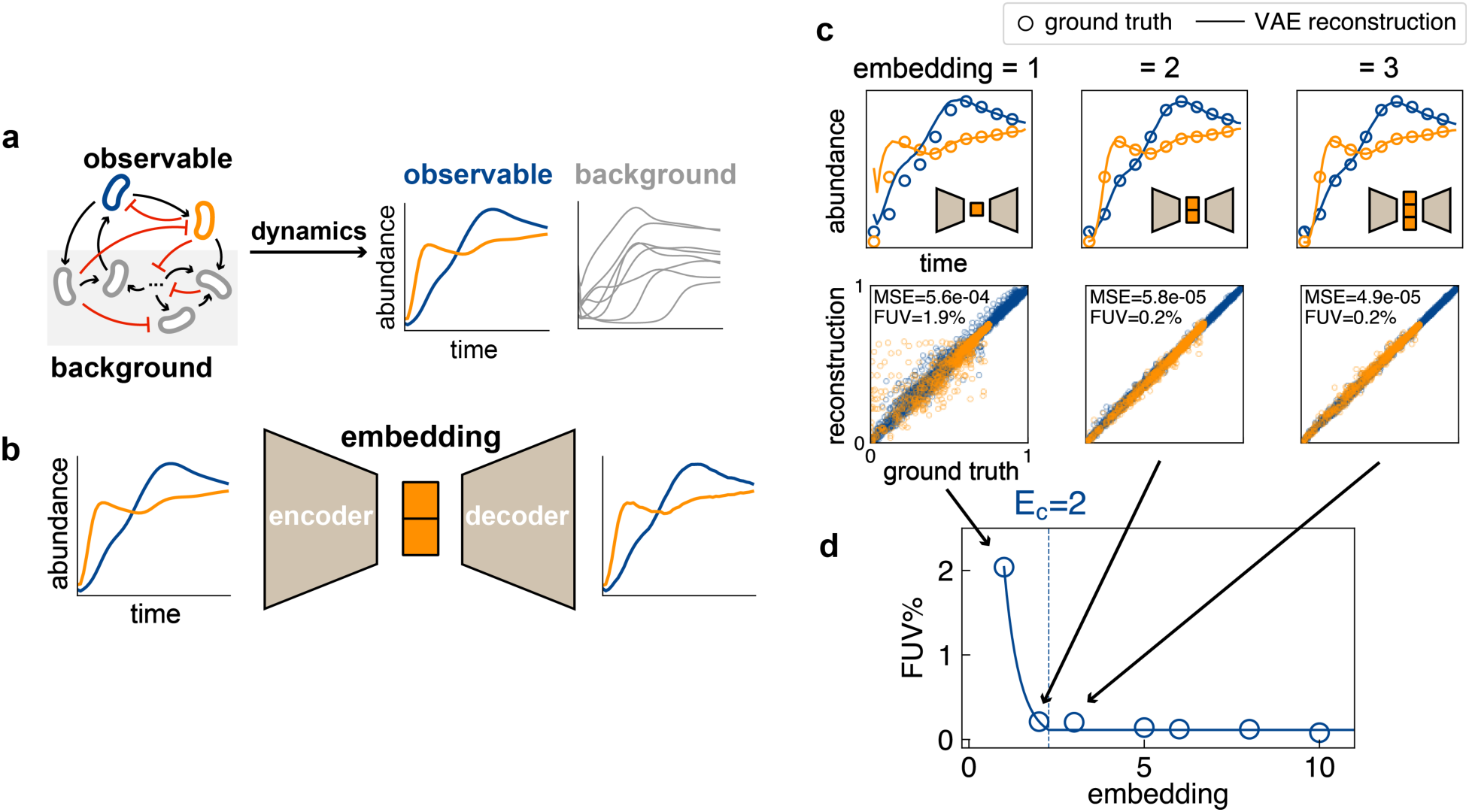
Variational autoencoder (VAE) reveals a low-dimensional representation of observed population dynamics. a) Observable microbial populations engage in complex ecological interactions with the background microbiota, giving rise to various temporal behavior. b) VAE is trained to compress the time series of the observables into latent space and reconstruct the dynamics from embedded representation. c) Training multiple VAEs with different embedding dimensions allows assessment of reconstruction quality across various compression levels. d) The critical dimension (*Ec*) is defined as the minimal embedding dimension that achieves a sufficient reconstruction quality, as measured by the fraction of unexplained variance (FUV). For the two observables in this community, an *Ec* of 2 suffices to perform optimal compression and reconstruction.

We define the critical dimension *Ec* as the smallest number of variables that preserve enough information to reconstruct the observed time series with high fidelity. To determine *Ec*, we used VAEs as a dimension reduction tool for the observed time series without extensive assumptions about the system (**Methods**). A VAE consists of an encoder and a decoder. The encoder compresses the input trajectories into a low-dimensional latent representation (embedding), and the decoder reconstructs the original data from this embedding (**Fig. 1b**). We trained a series of VAEs that differ in their embedding dimensionality but are otherwise structurally identical. Each model was trained solely on the observables, without access to background dynamics, which allowed us to determine the how the embedding dimension affects reconstruction quality (**Fig. 1c**).

We quantified reconstruction quality by using the fraction of unexplained variance (FUV) of the testing dataset, which is the mean squared error (MSE) between the ground truth and reconstructed time series, normalized by the total variance in the dataset. Increasing embedding dimension improved the reconstruction quality until reaching a threshold beyond which the performance plateaued (**Fig. 1d**). Embeddings larger than this threshold provided redundancy, while those smaller than the threshold caused information loss. We thus defined this threshold as *Ec* and extracted it by fitting a threshold exponential equation to the FUV-embedding curve (**Methods**). The example shown in **Fig. 1d** yielded a critical dimension of *Ec* = 2.

### Critical embedding dimension scales linearly with the number of observables

To determine *Ec* of the observable dynamics, we generated two types of simulation data. In the “fixed” case, simulations of the gLV model were performed with varying initial abundances of the observables and fixed initial background abundances (**Fig. 2a, Fig. S2a**). For the “random” case, the initial abundances of both observable and background populations were randomized across simulations (**Fig. 2b, Fig. S2b**).

**Figure 2.**
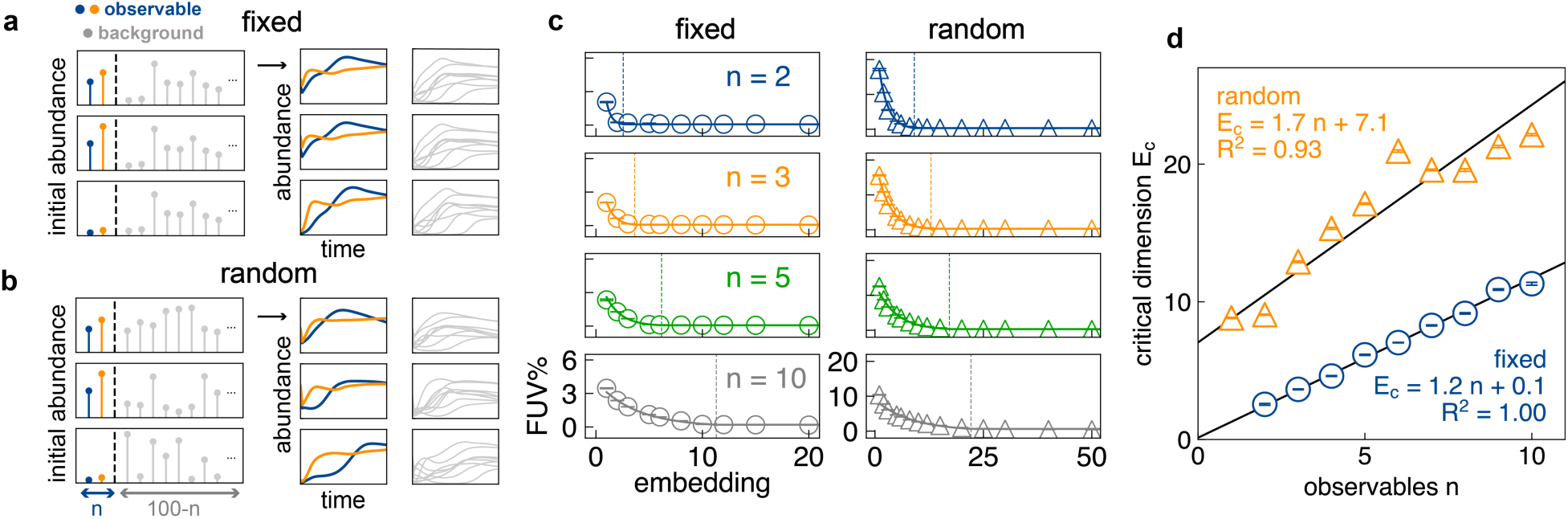
Critical dimension *Ec* scales linearly with the number of observables. a) “Fixed” background simulation: The initial abundances of observables vary; the initial abundances of background populations are fixed at non-equilibrium values across 10,000 simulations. b) “Random” background simulation: Initial abundances of all community members (observables and background) are randomized across 10,000 simulations. c) Assessment of the reconstruction quality (FUV, mean ± SE, 5 training replicates). Two columns correspond to fixed-background and random-background simulations. *n* denotes the number of observables. A threshold exponential curve is fitted to the data to derive the critical dimension *Ec*. *Ec* represents the minimum VAE dimension that allows sufficient reconstruction. Only *n* = 2, 3, 5, and 10 are shown. The rest of the FUV – embedding relationships can be found in **Fig. S3**. d) *Ec* (mean ± SE) exhibits robust linear scaling with the number of observables *n* for both fixed (blue, σ-weighted least-squares regression. Slope 1.157 ± 0.006 SE, *p* = 4×10^-14^; weighted R^2^ = 0.996) and random (orange, σ-weighted least-squares regression. Slope 1.725 ± 0.006 SE, *p* < 1×10^-16^; weighted R^2^ = 0.925) cases. This scaling indicates intrinsic low dimensionality of the simulated community dynamics.

For the “fixed” case, we simulated *n* = 2 to 10 observables for a given community. *N* = 1 was excluded because the FUV was already near zero at *Ec* = 1, making it infeasible to estimate a turning point from the FUV–embedding curve. For each choice of *n*, we extracted the corresponding *Ec* by training VAE models with different embedding dimensions to reconstruct the observed dynamics (**Fig. 2c**, left; three randomized communities with different parameter sets are shown in **Fig. S3a**). In each case, reconstruction quality improved with increasing embedding dimension and plateaued at *Ec* (corresponding to FUV ∼ 0).

We found that *Ec* scaled linearly with *n*: *Ec* = 1.2 *n +* 0.1 (**Fig. 2d; Fig. S3b**), which approximates the theoretical expectation *Ec* = *n*. When the initial abundances of the background populations are fixed, the time series generated by the same dynamical model are entirely governed by the initial abundances of the *n* observables. In this scenario, the ideal decoder is the original ODE model, which can generate the observable time-series solely based on these initial values. That is, the VAE could in theory learn to approximate the original ODE model.

We next examined the “random” case (**Fig. 2b, Fig. S2b**). Here, the full system has *N* = 100 degrees of freedom. Still, we found that *Ec* followed a linear correlation with *n*: *Ec* = 1.7*n +* 7.1 (*n* ranging from 1 to 10, **Fig. 2d**; three randomized communities with different parameter sets are shown in **Fig. S3c**). Due to the randomized background, *Ec* was consistently larger than the “fixed” case for each *n*. However, *Ec* was much smaller than *N*, underscoring the low dimensionality of observed time series, despite the much higher dimensionality of the full system. Our results are not limited to the use of VAE, but hold for other dimension estimation techniques, including Levina-Bickel algorithm^46^ (**Fig. S4**) and principal component analysis (**Fig. S5**). Moreover, the scaling law holds for communities with smaller sizes (10-member community, **Fig. S2c, Fig. S6 a&b**) and stronger interactions (**Fig. S2d, Fig. S6 c&d**).

In principle, for a gLV model with fixed parameters, the observable dynamics are solely governed by the initial abundances of the community. As the critical embedding captures the key elements describing the observable dynamics, we hypothesize the initial abundances can be projected onto the critical embedding to generate observable dynamics, enabling mechanism-free modelling of the observables. To test this notion, we followed a previously developed model pipeline^44^, by constructing MLP models that map the community initial conditions to the critical embedding, which was further decoded by the decoder component of the VAE to predict the observable time series (**Methods**). During training, we froze the parameters in the decoder, and trained the MLP models to minimize the MSE between predicted and ground truth observable time series. This combined MLP-VAE model enabled predict the observable dynamics with average R^2^ = 0.990 for 10-member communities (across 5 training replicates, 3 randomized communities, and observables *n* = 2, 3, 5, 8, 10), and R^2^ = 0.973 for 100- member communities (across 5 training replicates, 3 randomized communities, and observables *n* = 2, 3, 5, 8, 10) (**Fig. S7**), manifesting the potential applicability of this pipeline for data-driven community dynamics modelling, and confirming the VAE did not merely learn to memorize the samples in the training data.

The linear increase of the critical dimension *Ec* with *n* aligns with the analytic predictions by the dynamical mean-field theory of gLV^47–49^. When pairwise interactions are weak and randomly distributed, and the ecosystem is large (*N* → ∞), the background community becomes weakly coupled to a focal species only as a function of community biomass. In specific, when the system has a single fixed point and in the large *N* limit, community biomass becomes independent of the community’s initial conditions through averaging, giving rise to a slope-one linear relationship between the intrinsic dimension and the number of observables *n*. Importantly, the original theory assumes the gLV form, large ecosystem, and weak interaction limit. Our data-driven analysis robustly reproduces the linear scaling in the empirical regimes that depart significantly from those idealized assumptions.

### Linear scaling is generalizable in different simulated microbial ecosystems

To test the generality of our observations further, we explored the compressibility of additional microbial ecological scenarios through modelling (**Methods, Fig. S8**). We examined the following three models:

#### (1) Spatially structured communities with patch-dependent interactions^50^ (Fig. 3a).

We consider the dynamics of *n* focal species within a meta-community of 100 species distributed across 900 microbial patches on a 30 by 30 lattice grid. Different patches engage in distance-dependent interactions with one another. We consider the scenario when the background community has a fixed spatial organization while the focal species are randomly distributed across the patches. When applying VAEs to determine the critical dimension, we take the global abundances of the focal species as the observables, ignoring their spatial distributions.

1. (2) **Microbial communities with continuous immigration** (**Fig. 3b**). We simulated 100-member community dynamics subject to constant immigration from an external species pool^15^ for both “fixed” and “random” cases.
2. (3) **Horizontal transfer of plasmids within microbial communities** (**Fig. 3c**). We used a model^51^ describing the horizontal transfer of multiple plasmids within communities, with strain-level resolution. We simulated 100-member community transferring 5 different plasmids. During the simulation, we randomized the distribution of *n* target plasmids (*n* = 2, 3, 4, 5) across the 100 host microbial members while fixing the initial abundances of the 100 members and the 5-*n* background plasmids. We considered the total abundances of the target plasmids as the observables.

**Figure 3.**
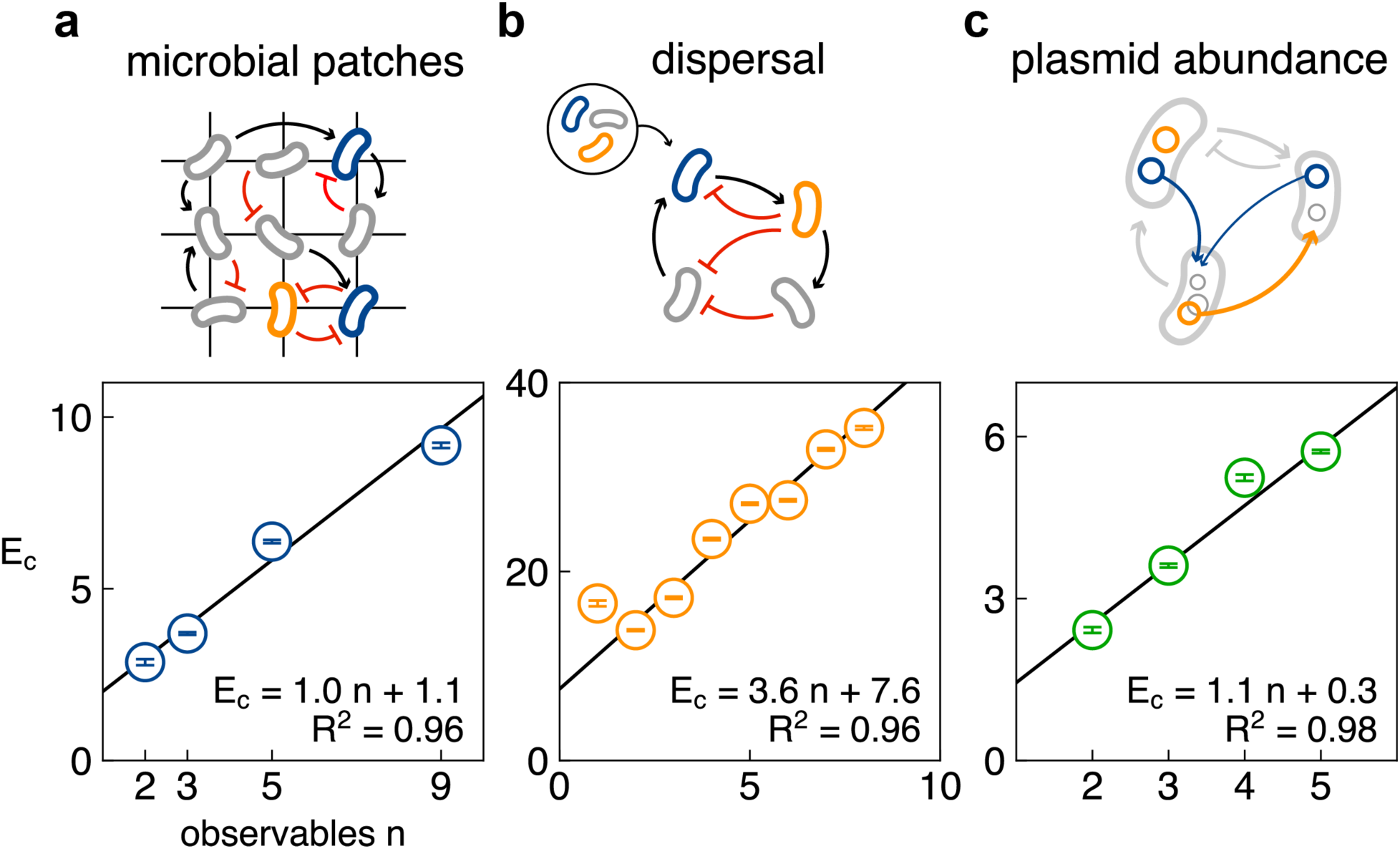
Linear scaling of *Ec* is generalizable to different simulated microbial ecological scenarios. a) Spatially structured microbial communities with patch-dependent interactions. *Ec* (mean ± SE) linearly scales with *n* (σ-weighted least-squares regression. Slope 0.955 ± 0.013 SE, *p* = 2×10^-4^; weighted R^2^ = 0.963). b) Microbial communities with continuous immigration (constant dispersal). *Ec* (mean ± SE) linearly scales with *n* (σ-weighted least-squares regression. Slope 3.57 ± 0.023 SE, *p* = 5×10^-12^; weighted R^2^ = 0.958). c) Horizontal transfer of plasmids within communities. The target plasmids (observables) are shown in blue and orange; the background variables are shown in grey. *Ec* (mean ± SE) linearly scales with *n* (σ-weighted least-squares regression. Slope 1.094 ± 0.018 SE, *p* = 3×10^-4^; weighted R^2^ = 0.976).

We quantified the critical dimension of the observed dynamics for each scenario with three sets of randomized parameters (**Methods**). For these ecosystems, the models’ degrees of freedom are drastically larger than the number of observables *n* (900*n* for microbial patches, 100 for communities with dispersal, and 100*n* for plasmid dynamics). Across the different simulations, *Ec* linearly scales with *n* (**Fig. 3, Fig. S9&S10**). For microbial patchy dynamics and plasmid dynamics, *Ec* are below 20 (**Fig. 3** **a&c, Fig. S9**), highlighting the simplicity in microbial spatial and plasmid dynamics. Indeed, models that ignore the spatial dimension^21,22^ have been used to describe gut microbiome dynamics, despite its spatial heterogeneity^52,53^. Plasmid abundance is less variable than community composition at high transfer rate^54^, and can be predicted by a coarse-grained metric of the community^51^. Our results provide a *post hoc* explanation why such simplifications can work. For communities with constant dispersal (**Fig. 3b, Fig. S10**), *Ec* is much larger (*Ec* = 3.6 *n* + 7.6), which may stem from the complex temporal dynamics the model was able to generate^15^.

### Linear scaling was applicable to experimentally assembled synthetic communities

To experimentally test the scaling law emerging from simulated data, we constructed synthetic communities each consisting of 62 engineered *E. coli* strains derived from the Keio collection^55^. Each strain was labeled with a unique 28-bp barcode on the chromosome through CRISPR-guided integration^56^ (**Methods, Table S1**). We randomly selected 4 strains as the carrier of 4 non-conjugative plasmids, each encoding a distinct fluorescent protein (mEGFP, mTagBFP2, LSSmOrange, and mCherry). The four fluorescent readouts, and optical density at 600 nm (OD), were taken as the five observables of the community.

We assembled communities with highly variable initial compositions at high throughput by strong dilution of a master starter mixture with a fixed composition (**Fig. 4a, Methods**). Assuming the cell number for each strain in each community approximately follows Poisson distribution^8^, we can estimate the average initial cell number was ∼ 0.5 – 1 for each background strain, leading to coefficient of variance of 100% (=1/1^0.5^) - 141% (= 1/0.5^0.5^), and ∼ 1.3 – 2.6 for each plasmid carrier, with standard deviation of 62% (=1/2.6^0.5^) - 88% (=1/1.3^0.5^) (**Methods**). Thus, the initial community compositions were highly randomized, corresponding to the “random” scenario. As a result, we tracked the fluorescence and OD dynamics of 887 communities for 48 hours with high-temporal resolution using microplate readers.

**Figure 4.**
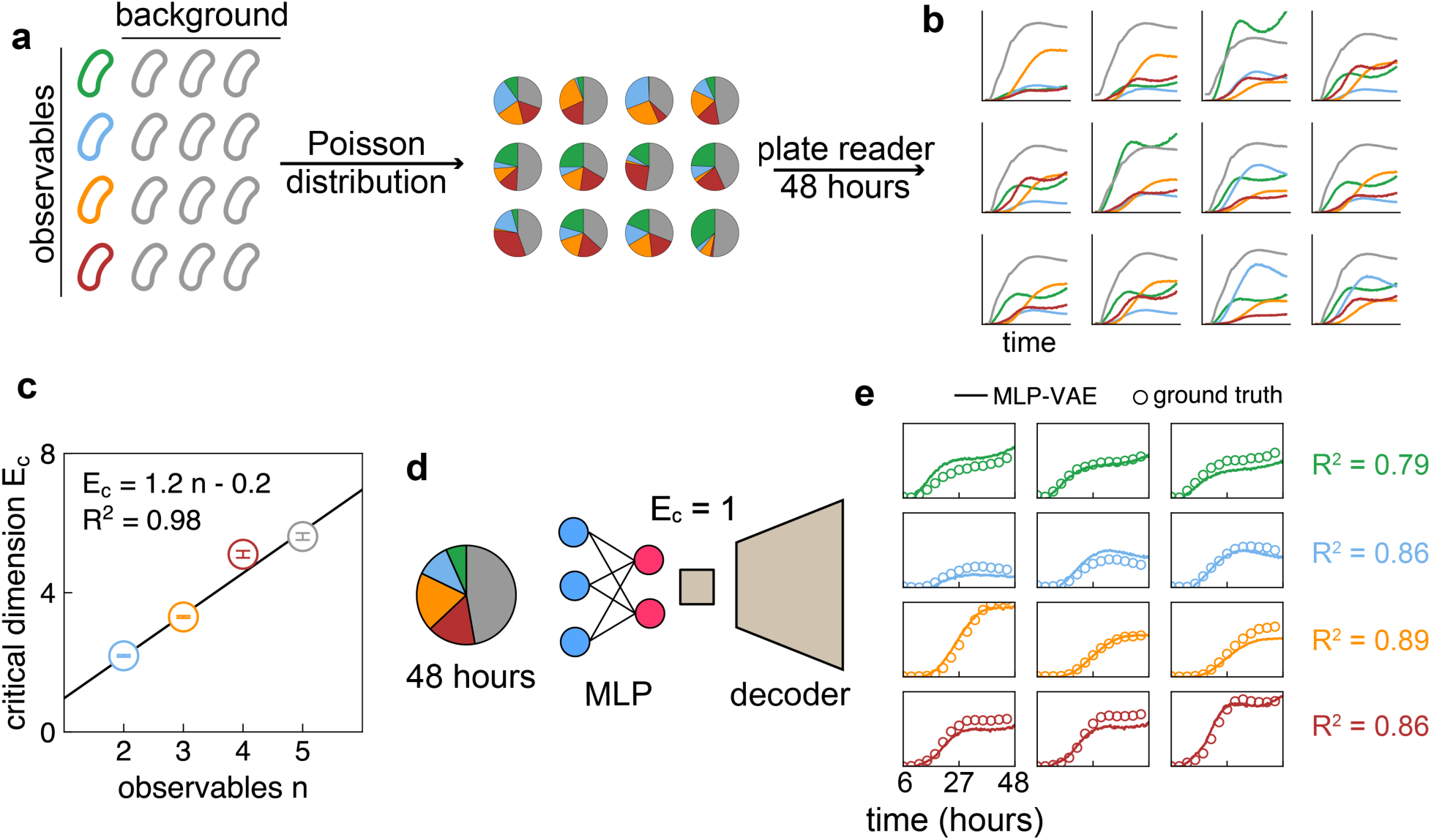
Experimental validation of the linear scaling in synthetic microbial communities. a) 62 uniquely barcoded *E. coli* strains were assembled into communities under limiting dilution, leading to random initial abundances. Four strains each carried a plasmid encoding one fluorescence protein (FP): mEGFP, mTagBFP2, LSSmOrange and mCherry. At time 0, the initial seeding cells for each strain approximately follow Poisson distribution, making the assembly of each community random. The communities were cultured in 384-well plates and measured for their fluorescence and OD600 for 48 hours. b) The observed dynamics (fluorescence and OD) exhibited rich temporal variability. c) *Ec* (mean ± SE) scales linearly with the number of observables, *n* (σ-weighted least- squares regression. slope 1.20 ± 0.03 SE, *p* = 5×10^-4^; weighted R^2^ = 0.982). Here *n* = 2 corresponds to mEGFP and mTagBFP2, *n* = 3 includes LSSmOrange, *n* = 4 includes mCherry, and *n* = 5 includes OD. d) Using multilayer perceptron (MLP) models to infer individual fluorescence dynamics based on endpoint plasmid-carrying strain abundances from NGS. For the four FPs, MLP model was attached to the decoder of the FP-specific VAE model with *Ec* = 1, and trained to map the relative abundance of the four plasmid0carrying strains at endpoint to the latent embedding, which was further used by the decoder model to infer the FP dynamics. e) The MLP-VAE model inferred the FP dynamics with high accuracy: R^2^ = 0.787 ± 0.010 for mEGFP, 0.862 ± 0.002 for mTagBFP2, 0.886 ± 0.006 for LSSmOrange, and 0.861 ± 0.003 for mCherry (mean ± SE over 10 independent 10-fold cross- validation trials).

These communities showed varied and rich temporal dynamics (**Fig. 4b**). We trained VAEs to determine *Ec* of *n* observables (*n* = 2, 3, 4, 5, **Fig. S11**), again revealing a linear scaling: *Ec* = 1.2*n* – 0.2 (**Fig. 4c**). The *Ec* values were much smaller than the strain pool size (*N* = 62).

Based on the linear scaling, we can estimate the critical dimension for each observable is approximately 1. Indeed, we found VAE models with *Ec* = 1 trained separately on each individual observable was able to reconstruct their dynamics with high fidelity (R^2^ = 0.982 for mEGFP, 0.972 for mTagBFP2, 0.990 for LSSmOrange, 0.984 for mCherry, and 0.973 for OD).

Measurement capability limits the sampling frequency of a microbiome to days, while bacteria growth and death operates on the scale of minutes to hours. This discrepancy results in the challenge of inferring bacterial growth dynamics based on snapshot sequencing measurements^57^. Here, as a proof of principle, we demonstrate the use of the endpoint abundances of the 4 plasmid-carrying strains to infer the fluorescence dynamics. We first quantified the endpoint composition of 289 communities through next-generation sequencing (**Methods**). We then constructed an MLP model with 10-fold cross-validation to map the endpoint abundances of the 4 plasmid-carrying strains to the *Ec* = 1 -dimensional embedding of each observable, based on which the decoder of the VAE will reconstruct the observable dynamics (**Fig. 4d, Fig. S12**). The MLP models predicted the temporal dynamics of the four observables with R^2^ = 0.79 for mEGFP, 0.86 for mTagBFP2, 0.89 for LSSmOrange, and 0.86 for mCherry.

### The VAE embedding can forecast target microbial dynamics and community compositional shifts

Microbiome composition can drastically change over time, exhibiting oscillations^10,15,16^ and transitions between alternative steady states^14^. The ability to forecast microbial dynamics and community compositional shift can be applied to microbiome engineering^58^ and ecological health surveillance^10,16^. Since the VAE embedding contains key dynamical information of the observables and the relevant background populations, we hypothesize it can be used to forecast target microbial dynamics and community compositional shifts. We used a previously published dataset with reported abrupt compositional shifts^14,59^. A soil microbiome and a pond water microbiome were inoculated separately into three different media (medium A: oatmeal broth; medium B: oatmeal + peptone; and medium C: peptone) and cultured for 110 days. Eight replicates were passaged in parallel for each of the six communities (Soil-A – C, and Water-A – C). For each community, the eight replicates together harbor 28 - 144 members (amplicon sequencing variants, ASV), with heterogeneity across replicates. During the 110 days, the communities exhibited different extent of compositional shifts^14^ (**Fig. S13**).

We first set out to determine whether the linear scaling exists between *Ec* and *n* for these communities. Due to the small data size for VAE training, we augmented the dataset through segmentation^60,61^ and combination^62–64^ (**Fig. 5a, Methods**). Briefly, we split the entire time series (of 80 or 110 days) into 30-day segments, with each segment differing from one another by a minimum of 1 day. We assumed that during each measurement, we could only observe the dynamics of randomly selected *n* members within the community. We emulated this process by randomly assembling the segmented time series of *n* members from the community as one datapoint to generate the dataset. Since this is a combinatorial process, the potential dataset size depends on both community richness and number of observables. Water-C community was low in richness, resulting in small dataset sizes, so we excluded it for our analyses. Our method yields datasets with sizes between 3,000 and 120,000 for the five communities of *n* = 1 – 5.

**Figure 5.**
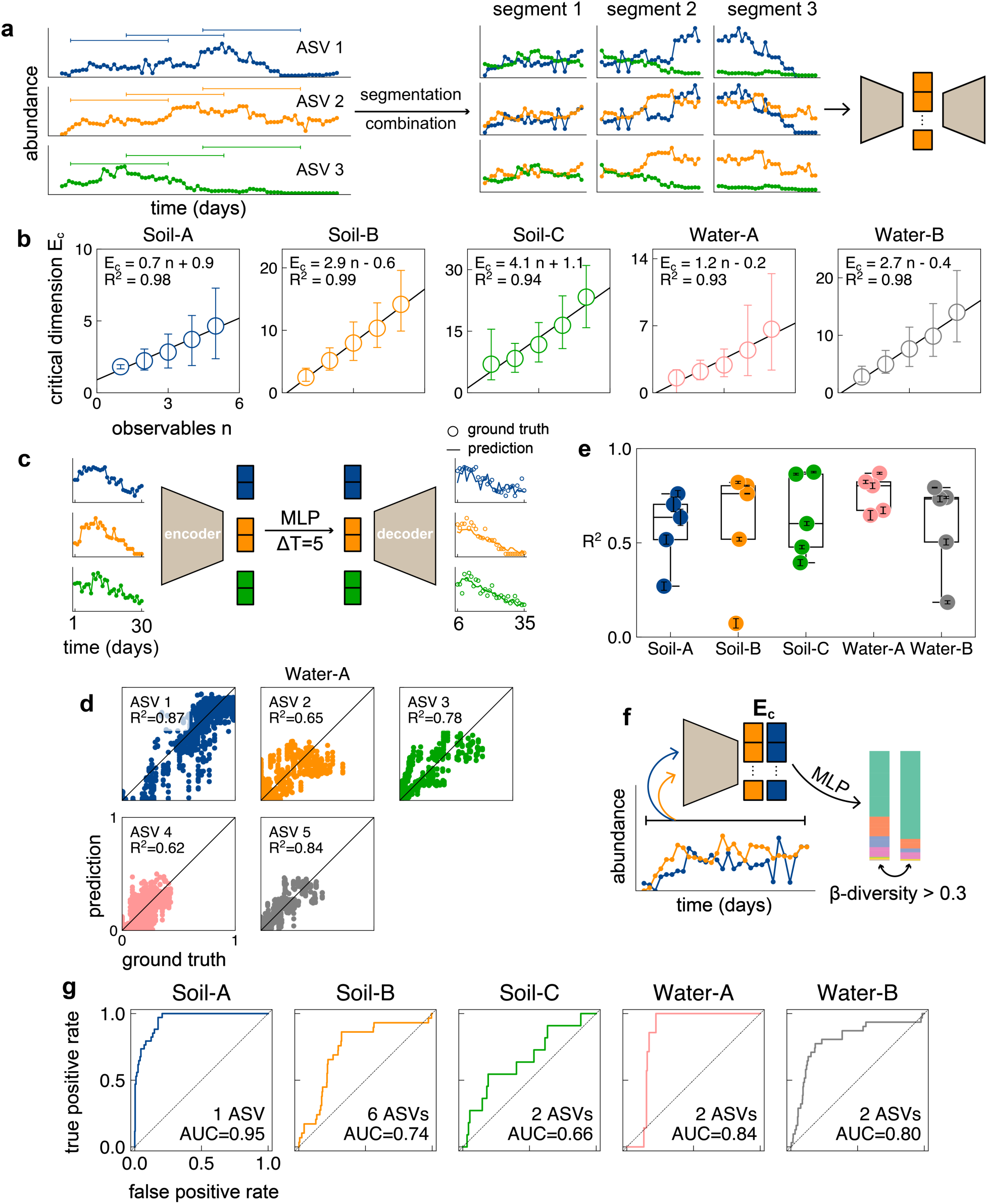
VAE embeddings enable forecasting of target dynamics and community composition shift. a) Time series of ASV data (80 or 110 days) were segmented into 30-day windows, and randomly combined to create augmented datasets of *n* ASVs for VAE training. b) *Ec* scales linearly with the number of observables, *n* across the five communities (Soil-A: Slope: 0.72, 95% confidence interval (CI) = [0.21, 1.28], bootstrap *p* = 4×10^- 3^; R^2^ = 0.975. Soil-B: Slope:2.86, 95% CI = [1.90, 4.02], *p* < 1×10^-3^; R^2^ = 0.994. Soil- C: Slope: 4.08, 95% CI = [1.92, 6.14], *p* = 2×10^-3^; R^2^ = 0.939. Water-A: Slope: 1.24, 95% CI = [0.19, 2.62], *p* = 8×10^-3^; R^2^ = 0.931. Water-B: Slope: 2.74, 95% CI = [1.48, 4.27], *p* < 1×10^-3^; R^2^ = 0.984.). *Ec* is defined as the embedding dimension that accounts for 90% of the data variance (FUV = 10%). c) An MLP model trained by using the VAE embeddings enabled forecasting of future ASV abundances 5 days ahead. It maps the embedding of ASV dynamics during days 1 – 30 to the embedding of ASV dynamics during days 6 – 35. Open circles: measured relative abundances; curves: predicted dynamics. d) Example prediction of the top five prevalent ASVs in the Water-A community. The MLP model forecasts the next five days dynamics with R^2^ ranging from 0.62 to 0.87. e) Summary of MLP prediction performance across communities. Median R^2^ over five replicates: Soil-A: 0.64, Soil-B: 0.76, Soil-C: 0.60, Water-A: 0.80, and Water-B: 0.73. For each ASV, mean ± SD R^2^ over five training replicates is shown. f) The segmented time series (day 1 – 30) were encoded into the embedding of a VAE trained on 1-target time series. The resulting embeddings of the selected ASVs were stacked and used to predict the occurrence of large community shift (Bray-Curtis β- diversity > 0.3) on the 31st day by an MLP model. g) Receiver Operating Characteristic curve analyses demonstrate strong predictive power on large community shifts using the stacked embeddings. The model achieved area under the curve (AUC) of 0.96 for Soil-A, 0.74 for Soil-B, 0.60 for Soil- C, 0.84 for Water-A, and 0.82 for Water-B. Results shown are from one representative training replicate.

To prevent data leakage, we performed 8-fold cross-validation based on the 8 replicates during the training of the VAE models, where in each cross-validation fold, we took one replicate community as the test dataset, and the other 7 as train dataset. The average FUV did not follow the threshold exponential dependence on the embedding dimension (**Fig. S14a**), potentially attributed to the experimental noise the model could learn at high embedding dimension. Thus, we estimate *Ec* as the embedding dimension that explains 90% of data variance (FUV = 10%). Across the five communities, we found *Ec* to increase with *n* linearly (**Fig. 5b**), though the slope and intercept of the linear relationship varies across these communities. As we show in **Fig. S14b**, the linear scaling is robust against the specific choice of the threshold FUV. Numerically, we confirmed the estimated *Ec* of the augmented dataset is similar to the *Ec* of a specific choice of *n* observables, based on gLV simulations under the “random” case (**Fig. S15**).

The linear correlation between *Ec* and *n* suggests each individual population can be embedded into the latent space with the same dimension. Given this notion, a VAE model that embeds the individual time series of all ASVs offers the most flexibility, where one can stack the embedding of the observables for downstream tasks. We trained VAEs with *Ec* = 3 for Soil-A, 5 for Soil-B, 7 for Soil-C, 2 for Water-A, and 5 for Water-B communities based on all the available data across the 8 replicates. We use the embeddings from these VAEs for the downstream tasks for target dynamics and community shift forecasting.

The critical embedding of the observables records information that explains 90% of the variance, filters out additional noise, and reduces the input dimension for constructing downstream regression models. Previous works have shown observed time series can be used to forecast its future dynamics^14^ through delay embedding^59^, sequential locally weighted global linear maps^65^, and autoencoder embedding. For each community, we trained an MLP model that operates in the VAE’s latent space, projecting the critical embedding of a previous time segment (day 1 to day 30) to the critical embedding of a latter segment (day 1+ ΔT to day 30 + ΔT) (**Fig. 5c**). We trained the MLP models based on the dynamics of the five most prevalent ASVs in each community, and performed 8-fold cross-validation based on the 8 replicates. For the Water-A community, the abundances of the next ΔT = 5 days can be predicted with R^2^ of 0.62 – 0.87 (**Fig. 5d**) for the top five ASVs. Across the five communities, the median prediction accuracy ranges between R^2^ = 0.60 to 0.80 (**Fig. 5e, Fig S16 a-e**). With prediction accuracy drops with increasing ΔT (**Fig. S16f**).

During abrupt shift, the abundances of the dominant members vary drastically.

Since the embeddings can forecast the dominant ASV’s future abundance, we hypothesize it can also forecast the shift in community composition. We quantified the extent of community shift by the Bray-Curtis β-diversity before and after a given time point (**Methods**), consistent with the original study^14^. We stacked the critical embeddings of the most prevalent 1 – 10 ASVs and used them as inputs for MLP models to predict community shift at the endpoint the time series segment (between day 30 and day 31) (**Fig. 5f**). To balance the dataset, we used weighted binary cross entropy as the loss function for training and ensured no data leakage by training with 8-fold replicate-based cross-validation. While we found the MLP models were not able to perform regression on the actual value of β-diversity well, the critical embeddings contain sufficient information to forecast whether β-diversity will exceed a threshold value (0.3 or 0.2) at the endpoint of the segment (**Fig. 5f, Fig. S17**). For Water -A and -B, and Soil -A and -B communities, the critical embeddings could forecast large compositional shifts with area under the Receiver Operating curve (AUC) between 0.74 and 0.96 (**Fig. 5g**), while in Soil-C community achieved AUC = 0.6. The Receiver Operating Characteristic curves based on the embedding of 1 – 10 ASVs are shown in **Fig. S17**.

### Human vaginal microbiome dynamics is low-dimensional

Dynamics of natural communities often respond to changes in environmental conditions. For instance, vaginal microbiome is also subject to drastic compositional shifts during menstruation^10^, and are subject to dysbiosis like bacterial vaginosis (BV) ^16^. To test whether the same linear scaling exists in natural communities with environmental fluctuations, we used a published dataset on human vaginal microbiome^16^, sampled for 70 days. Among the 25 women in the study, 15 experienced symptomatic bacterial vaginosis, 6 were diagnosed with asymptomatic BV, and 4 were healthy. We similarly augmented the dataset as in the previous section, except we split the time series into 14-day segments, yielding dataset sizes of 15,000 and 52,000 for *n* = 1 - 5. By performing 24-fold cross-validation during VAE training (**Fig. S18a**, excluding one subject for low richness in the microbiome), we again observed linear scaling between *Ec* and *n* across different FUV thresholds, with *Ec* = 2.9 *n* + 0.3 for FUV = 10% (**Fig. S18b**).

Previously, Bashan et al. showed that human gut and mouth microbiomes are governed by ecological dynamics universal across individual hosts, while skin microbiomes are not^66^. Here, we observed the vaginal microbiome dynamics across subjects can be embedded into the same low-dimensional latent space. This observation suggests universal low-dimensional dynamics also exists for human vaginal microbiome.

## Discussion

In this work we demonstrated a data-driven approach to quantify the critical dimension to represent observable microbial community dynamics. Repeatedly, we found the critical dimension scales linearly with the number of observables. This linear scaling is robust against variations in community size, interaction types, and interaction complexity, indicating an intrinsic simplicity of microbial temporal dynamics. Our use of VAE for dimension reduction focuses on revealing an intrinsic property of a microbial community as a dynamical system. This focus differs from the typical application of dimensionality reduction^67^, which often focuses on feature extraction for specific downstream applications. While VAEs are not unique in revealing this scaling property (**Figs S4 & S5**), their advantage is the ability to provide an unbiased embedding with quantifiable reconstruction quality, thus offering a flexible and rigorous framework for representing dynamical systems.

Past studies have demonstrated different types of simplicity in microbial communities, highlighting the convergence in community compositional^68^ and functional structures^69^, or predictive power of simple theoretical models^70–76^, usually at a snapshot or at equilibrium. Some of these simplicities were found to be driven by metabolic functional convergence^68,77^, and growth competition^78^, while others were hypothesized to be due to large-number averaging^75^, time scale decoupling^57^, or system-specific mechanisms^70^. In contrast, here we define and quantify a general form of simplicity: *the intrinsic low dimensionality of the microbial dynamics, irrespective of the nature of interactions*. This property underscores that complexity at a mechanistic level does not necessarily translate to complexity at the dynamical system level. Our results suggest the feasibility of prediction and control over the target microbes without knowledge of the background communities, opening doors to new approaches for microbiome prediction and engineering.

Coarse graining is an extensively adopted practice to reduce the number of variables of a microbial community model and accelerate model fitting and simulation. Due to limited measurement resolution, coarse graining becomes a necessity to derive well-constrained models based on reliable measurements. Researchers adopted *ad hoc* strategies like grouping of individual members^18,38,79^ and removal of non-dominant taxa^18,21,22^ to generate coarse-grained gLV models. Similar approaches have been adopted to model synthetic gene circuits ^80–82^ while ignoring the background host gene regulatory network and physiology^83^, and viral infection dynamics while ignoring the complex immune system^84–86^. Our results highlight the low dimensionality of a focal population in comparison to the entire community, suggesting the theoretical feasibility of such simplifying approach. Moreover, the VAE embedding offers a platform for constructing coarse-grained models with minimum assumptions. A given set of information of the system can be mapped to the embedding through supervised learning (e.g. **Fig. 4, 5, S7**), enabling evaluation of appropriate coarse-graining, *post hoc* mechanistic interpretation, and hypothesis generation.

The deduction of the embedding using VAE requires sufficient data for training. Currently, microbiome datasets with high temporal resolution are generally lacking, despite containing rich information on the system’s dynamics and stability^10,14^. High- throughput automation of the experimental pipeline^12,87–89^ could greatly facilitate the data generation. Moreover, our results suggest label-based, cost-effective quantifications can yield important insights into the entire system, without the need for sequence-based profiling the entire community.

An alternative to circumvent the data requirement for a specific experimental system is to leverage pre-trained foundation models^90^. These models could facilitate the analyses of experimental data that’s relatively scarce, by incorporating learned knowledge based on published datasets from diverse sources. In our results, the linear scaling between the critical dimension and the number of observables suggests the time series of one microbe can be embedded in a constant dimension. Therefore, a community dynamics foundation model should be designed to handle individual microbial time series, maximizing its analytical flexibility to adapt to differently sized microbiomes.

In summary, our work used neural network models to represent observable microbial temporal dynamics. The expressivity of the neural network permits the delineation of independent variables, as well as the measurement of data complexity. The observed linear scaling between the critical dimension and the number of observables points to the simplicity in the structure of microbial community dynamics, providing insights into the coarse-grainability and organization principles underlying the microbial communities.

## Acknowledgments

We thank Ophelia Venturelli, Yili Qian, Jaron Thompson, Sarvesh Menon, Dongheon Lee, and Ashwini Shende for helpful discussions. We thank Irida Shyti for assistance with setting up the GitHub repository. We thank Duke Compute Cluster for resources on high performance computing. D.K. also acknowledges partial support of his time by the Office of the Under Secretary of Defense for Research and Engineering under award number FA9550-22-1-0379. This work was partially supported by DARPA (D.K., L.Y. HR0011-23-2-0008) and National Institute of Health (L.Y. R01AI125604, R01GM098642).

## Author contributions

Z.Z. and L.Y. conceived the research. Z.Z. and X.C. performed the experiments. Z.Z., E.Ş., and G.H. performed the numerical simulations of mathematical models. Z.Z. performed neural network training and analysis. Z.Z., Y.B. and L.Y. wrote the manuscript. All authors read and contributed to the manuscript.

## Declaration of generative AI and AI-assisted technologies in the writing process

During the preparation of this work the authors used ChatGPT to improve the language of the manuscript. After using this tool, the authors reviewed and edited the content as needed and take full responsibility for the content of the published article.

## Resource availability

### Materials availability

Genetic constructs generated for this study are available upon request.

### Lead contact

Further information and requests for resources should be directed to and will be fulfilled by the lead contact, Lingchong You.

### Data and code availability

All original code has been deposited at You Lab GitHub Repository: https://github.com/youlab/Low_dimensional_observables. Simulation data, neural network models, and plate reader data have been deposited at Zenodo and are publicly available: https://zenodo.org/records/15659948.

