## Supplementary Materials for "Linear scaling reveals low-dimensional structure in observable microbial dynamics"

* Corresponding author

This PDF includes:

- Methods
- Figure S1 – S18
- Table S1

### Methods

#### 1. Computational methods

Numerical simulations of bounded gLV model and gLV with constant dispersal were performed using Matlab R2023a. Numerical simulations of microbial patch dynamics were performed using Matlab R2020a. Numerical simulations of plasmid dynamics were performed using Python 3.12.4. All machine learning analyses were performed using PyTorch 2.5.1. Specific versions of individual software can be found in the requirements.txt file at https://github.com/youlab/Low_dimensional_observables.

##### 1.1. Variational autoencoder (VAE) to estimate critical dimension

###### 1.1.1. Simulated data

We implemented 1D convolutional VAEs to learn the low-dimensional latent representation of multivariate time series data. We deployed multiple different variants of the VAE. The first one is used in the analyses of the following figures: (**Fig. 1 b – d; 2; 3 a & c; S3; S6 a & b; S9; S15**). The input consisted of *n* channels (corresponding to *n* observables), each with 50 time points.

- **Encoder:** The encoder comprised three 1D convolutional layers, each with 32 output channels, a kernel size of 3, stride of 1, and padding of 1, and two linear layers. Each convolutional layer was followed by a LeakyReLU activation. The output feature maps were flattened and passed through two separate linear layers (each with input size of 32 × 50, and output size of *E*) to compute the latent mean $\mu$ and log-variance $\log \sigma^2$ vectors, both of dimension *E* (embedding dimension).
- **Latent space:** We sampled latent vectors $z \sim \mathcal{N}(\mu, \sigma^2)$
- **Decoder:** The decoder first projected the latent vector back to a 32 × 50 - dimensional vector via a linear layer followed by LeakyReLU. This vector was reshaped and passed through three 1D transposed convolutional layers each

followed by a LeakyReLU activation, mirroring the encoder's structure in reverse. The final layer mapped the features back to *n* output channels, reconstructing the input shape.

- **Loss function:** The loss function for VAE to perform reconstruction $\boldsymbol{s}_t^{rec}$ from original data $\boldsymbol{s}_t^{ori}$ is defined by two parts. The first part is the mean square error (MSE) between original data and reconstructed time series to ensure reconstruction quality. The second part is a regularization term of the Kullbeck-Leibler divergence (with weight of $\alpha = 1\times10^{-4}$) that quantifies the discrepancy between the latent space distribution of the data and a normal Gaussian distribution:

$$KLD = -\frac{1}{2}\mathbb{E}[1 + \log(\sigma^2) - \mu^2 - \sigma^2]$$

$$Loss = L_{MSE}(\boldsymbol{s}_t^{ori}, \boldsymbol{s}_t^{rec}) + \alpha KLD$$

10,000 simulations were generated for each choice of *n* observables. 8,000 samples were used as the training data, and 2,000 as testing, based on which we assessed the FUV. Training was performed with Adam optimizer, learning rate of $1\times10^{-3}$, batch size of 64, for 100 epochs. Model at last epoch was used to assess the FUV. Five independent trainings (from five different initial weights) for each embedding dimension were performed.

For the bounded gLV model with stronger interactions (**Fig. S2d**, **Fig. S6 c&d**) and gLV model with constant dispersal (**Fig. 3b, Fig. S10**) and, four 1D convolutional layers with 64 channels were used for better model flexibility. Training was performed with Adam optimizer, learning rate of $1\times10^{-3}$, batch size of 64, for 20 epochs. Five independent trainings (from five different initial weights) for each embedding dimension were performed.

#### 1.1.2. Experimental data from engineered communities

For the data from engineered communities, the input consisted of *n* channels (corresponding to *n* observables), each with 168 time points. The VAE encoder consisted of three 1D convolutional layers, each with 32 output channels, a kernel size of 3, stride of 1, and padding of 1, followed by LeakyReLU activations. The resulting feature maps were flattened and passed through two fully connected layers to compute the latent mean and log-variance, together forming the complete encoder.

The decoder first projected the latent vector back to a 32 × 168 - dimensional vector via a linear layer followed by LeakyReLU. This vector was reshaped and passed through three 1D transposed convolutional layers each followed by a LeakyReLU activation, mirroring the encoder’s structure in reverse. The final layer mapped the features back to *n* output channels, reconstructing the input shape.

887 random communities were generated. 709 datapoint were used as the training data, and 178 as testing, based on which we assessed the FUV. Training was performed with Adam optimizer, learning rate of $1\times10^{-3}$, learning rate decay of 1% for every epoch, batch size of 16, for 200 epochs. Model at last epoch was used to assess the FUV. Five independent trainings (from five different initial weights) for each embedding dimension were performed.

#### 1.1.3. Community dynamics data from the literature

For the environment-derived communities, the input consisted of *n* channels (corresponding to *n* observables), each with 30 time points. For the human vaginal microbiome data, the input consisted of *n* channels (corresponding to *n* observables), each with 14 time points. The VAE encoder consisted of three 1D convolutional layers, each with 16 output channels, a kernel size of 3, stride of 1, and padding of 1, followed by LeakyReLU activations. The resulting feature maps were flattened and passed through two fully connected layers to compute the latent mean and log-variance, together forming the complete encoder.

The decoder first projected the latent vector back to a 16 × T (T = 14 or 30) -dimensional vector via a linear layer followed by LeakyReLU. This vector was reshaped and passed through three 1D transposed convolutional layers each followed by a LeakyReLu activation, mirroring the encoder's structure in reverse. The final layer mapped the features back to *n* output channels, reconstructing the input shape.

For the environment-derived communities, based on the data augmentation scheme, n = 1 gave 3,000 – 8,000 datapoints, and n = 5 gave 40,000 – 120,000 datapoints, for the five communities. 8-fold replicate-based cross-validation was performed for the training and evaluation of the VAE performance. Each fold of training was performed with Adam optimizer, learning rate of $5\times10^{-4}$, learning rate decay of 5% for every epoch, batch size of 64, for 30 epochs. Model at last epoch was used to assess the FUV. Three independent trainings (from three different initial weights) for each cross-validation fold and each embedding dimension were performed.

For human vaginal microbiome dataset, based on the data augmentation scheme, n = 1 gave 14,000 datapoints, and n = 5 gave 88,000 datapoints. 24-fold subject-based cross-validation was performed for the training and evaluation of the VAE performance. Each fold of training was performed with Adam optimizer, learning rate of $5\times10^{-4}$, learning rate decay of 5% for every epoch, batch size of 64, for 30 epochs. Model at last epoch was used to assess the FUV. Three independent trainings (from three different initial weights) for each cross-validation fold and each embedding dimension were performed.

##### 1.1.4. Estimating the critical dimension $E_c$ using threshold exponential function

For the data from numerical simulations and the synthetic community, to estimate the critical dimension, we fitted a threshold-exponential function to the FUV-embedding data:

$$y = \begin{cases} ce^{-b(x-E_c)}, & x \leq E_c \\ c, & x > E_c \end{cases}$$

To ensure the robustness of our conclusions, five VAEs were trained with different initial weights for each embedding dimension, the mean and standard error (SE) FUV were calculated across the five replicates. σ-weighted least-squares regression was performed using scipy.optimize.curve_fit function with input of the mean ± SE FUV. The estimate $E_c$ and standard error from the regression were obtained. Depending on the data, we chose the initial values for the fitting differently.

Linear regression between $E_c$ and $n$ was further performed to determine the linear scaling between the two variables. σ-weighted least-squares regression was performed using scipy.optimize.curve_fit function with input of the mean ± SE $E_c$. Two-sided t-test was performed to determine the statistical significance of slope is deviating from 0.

#### 1.1.5. Estimating the critical dimension $E_c$ in longitudinal microbiome datasets.

For the environment-derived communities and human vaginal microbiome, we estimated $E_c$ as the embedding dimension that explains $x$ percent of the total variance (FUV = 100-$x$). For each cross-validation fold, three VAEs with different initial weights were trained, and the average FUV was calculated. Then, based on the average FUV curves of each fold, 1,000 times of bootstrapping was performed. In each bootstrap iteration, we randomly sampled 8 (for environment-derived communities) or 24 (for human vaginal microbiome) FUV curves and computed the average FUV curve. We then estimated $E_c$ based on linear interpolation of this average FUV curve. Based on the

1,000-time bootstrapping, we estimated the 95% confidence interval (CI) of $E_c$ for a given *n*.

Based on the 1,000-time bootstrapping, we further performed linear regression on each bootstrap replicate, based on which we obtained the mean and 95 % CI for the slope and intercept. Two-sided empirical p-value was calculated for whether the slope is significantly deviating from 0: $p = 2 \times \min\left(P(slope \leq 0), (slope \geq 0)\right)$.

#### 1.2. Multilayer perceptron models (MLP)

In this work we constructed multiple MLP models to map between the critical embedding of observable time series to different objectives, or vice versa. For these downstream tasks, the first thing we did is to train a VAE model with a given critical dimension based on all the data available (train + test) before using the critical embedding for these downstream tasks. All the training are performed in the same way we obtained the $E_c$. During the training of the MLPs, the weights of the VAEs were frozen.

##### 1.2.1. MLP model for target dynamics modeling based on initial abundances

For 10-member communities (**Fig. S7 a&b**), the MLP model took as input the initial abundances of the 10 community members. The model consisted of three fully connected layers: the first two layers each with 32 neurons, and a final output layer with $E_c$ neurons to predict the latent embedding that was subsequently decoded by a pretrained VAE decoder to reconstruct the time series. The model was trained using 8,000 datapoints and evaluated on 2,000 held-out samples. Training was performed using the Adam optimizer with an initial learning rate of $1\times10^{-2}$, a learning rate decay of 1% per epoch, a batch size of 64, and for 200 epochs. The model was optimized using MSE between the predicted and ground-truth time series. Model at last epoch was evaluated for the performance. Five independent trainings (from five different initial weights) were performed.

For 100-member communities (**Fig. S7 c&d**), the MLP model took as input the initial abundances of the 100 community members. The model consisted of four fully connected layers: three layers each with 128 neurons, and a final output layer with $E_c$ neurons to predict the latent embedding that was subsequently decoded by a pretrained VAE decoder to reconstruct the time series. Due to convergence issues with 8,000 training samples, a larger dataset was generated, consisting of 80,000 samples for training and 20,000 for testing. Training was performed using the Adam optimizer with an initial learning rate of $2\times10^{-3}$, a learning rate decay of 1% per epoch, a batch size of 64, and for 100 epochs. The model was optimized using MSE between the predicted and ground-truth time series. Model at last epoch was evaluated for the performance. Five independent trainings (from five different initial weights) were performed.

#### 1.2.2. MLP model for FP dynamics inference

For the inference of the fluorescence dynamics based on endpoint plasmid carrier abundances (**Fig. 4e, Fig. S12**), the MLP model took as input the endpoint relative abundances of four community members. It consisted of four fully connected layers: first three layers each with 256 neurons, and a final output layer to predict the latent embedding of dimension $E_c = 1$, which was subsequently decoded by a pretrained VAE decoder to reconstruct the time series. Due to the limited dataset size ($n = 289$), 10-fold stratified cross-validation was performed, stratified by the maximum value in the time series with 5 bins. Training for each fold was performed using the Adam optimizer with an initial learning rate of $3\times10^{-4}$, a learning rate decay of 1% per epoch, a batch size of 16, and for 100 epochs. The model was optimized using MSE between the predicted and ground-truth time series. Model at last epoch was evaluated for the performance.

Ten independent trainings (from ten different initial weights) for each FP and each fold of cross-validation were performed.

#### 1.2.3. MLP model for target dynamics forecasting

For forecasting the target ASV dynamics (**Fig. 5 c-e, Fig. S16**), the MLP model operated in the latent space and took as input the critical embedding of an earlier time segment (days 1–30), extracted using the VAE encoder. The model consisted of seven fully connected layers: the first six layers each with 512 neurons, and a final output layer with $E_c$ neurons, predicting the embedding for a future segment (days 1 + ΔT to 30 + ΔT), which was then decoded by the VAE decoder to forecast the future time series. To avoid data leakage, 8-fold replicate-based cross-validation was performed. Training for each fold was carried out using the Adam optimizer with an initial learning rate of $1×10^{-3}$, a learning rate decay of 1% per epoch, a batch size of 32, and for 200 epochs. The model was optimized using MSE between the ground-truth time series (days 31–30 + ΔT) and the predicted time series. Model at last epoch was evaluated for the performance. Five independent trainings (from five different initial weights) for each fold of cross-validation were performed.

#### 1.2.4. MLP model for forecasting large community shifts

For forecasting the large community shifts (**Fig. 5 f&g, Fig. S17**), the MLP model took as input the stacked critical embeddings of the previous time segment (days 1–30), extracted from the VAE encodings of 1–10 ASVs. The model consisted of three fully connected layers: the first two layers each with 32 neurons, and a final output layer with one neuron, predicting the β-diversity between days 30 and 31 (see Section 1.5.1 for details on the metric calculation). To prevent data leakage, 8-fold replicate-based cross-validation was performed. Training for each fold was conducted using the Adam optimizer with an initial learning rate of $1×10^{-3}$, a learning rate decay of 2% per epoch, a batch size of 16, and for 40 epochs. The model was optimized using weighted binary cross-entropy loss with logits, to address class imbalance. Model at last epoch was evaluated for the performance. Five independent trainings (from five different initial weights) for each fold of cross-validation were performed.

### 1.3. Simulations of observable dynamics

#### 1.3.1. Bounded gLV model

For simulations in **Fig. 1, 2**, and **S1 – 7**, we used a gLV model that generates bounded dynamics of each member[1], with absolute abundance between 0 and 1.

$$\frac{d\boldsymbol{s}}{dt} = \boldsymbol{\mu s}(1 - \boldsymbol{s} - \frac{\sigma}{1 + \boldsymbol{\alpha s}} - \boldsymbol{\gamma s})$$

Here $\boldsymbol{s}$ is the abundance of the different members, $\boldsymbol{\mu}$ is the maximum growth rate, and $\sigma$ is the environmental stress imposed on the community. Different members interact positively with coefficients $\boldsymbol{\alpha}$ and negatively with coefficients $\boldsymbol{\gamma}$. The interaction matrices $\boldsymbol{\alpha}$ and $\boldsymbol{\gamma}$ has zero diagonal terms. The growth rate of each member follows a uniform distribution $\mu \sim U(0.5, 1)$. The interactions between two members follow uniform distribution $\gamma \sim U(0, 0.4)$ and $\alpha \sim U(0, 0.4)$. The environmental stress for each member follows a uniform distribution $\sigma \sim U(0.05, 0.15)$. For the communities with stronger interactions (Fig. S2d, Fig. S6 c&d)

Two types of simulations are performed using this model. The first type is performed by varying the initial abundances of the observables in every simulation, while fixing the initial abundances of the background across simulations. This is denoted as the “fixed” case in the main text. The second type of simulation is performed by randomizing the initial abundances of the entire community in every simulation, denoted as the “random” case. In both cases, the randomization of the initial abundances follows a uniform distribution of $s(0) \sim U(0, 0.2)$. ODE integration was performed for 300 arbitrary unit (a.u.), and observable abundances were recorded every 6 a.u., yield time series with length of 50. To avoid observing trivial dynamics (e.g., exponential decay), we randomly chose the observables from the ones with final abundances > 0.1 at the end of one simulation instance.

#### 1.3.2. Microbial patches

To simulate patchy microbial communities that engage in distance-dependent interactions (**Fig. 3 a, S8a, S9 a&b**), we consider a meta-community with 900 individual microbial patches each sitting on the vertex of a 30 by 30 lattice. In our simulation, we took a community size of *N* = 100. To assign the (*N*-*n*) background populations, we assume each microbial patch consists of one species, with a given initial abundance. Then, *n* focal species are randomly assigned onto the lattice with fixed locations but variable initial abundance across simulations. For two patches, $s_{ij}$ and $s_{mn}$ which sit on lattice point (i, j) and (m, n), we assume their interaction strength scales as the inverse of the Euclidean distance between these two patches, $\gamma_{ij,mn} = \gamma_0 / d_{ij,mn}$, where $\gamma_0$ stands for the interaction strength between the two species in well-mixed condition. Since certain grid points will have two species, the interaction strength between the two will be $\gamma_0$. Thus, for the same pair of microbes, their interaction when co-located is the same as when they are sitting on two adjacent grid points. We take the dynamics of the patches as follows:

$$\frac{d\boldsymbol{s}}{dt} = \boldsymbol{\mu s}(1 - \boldsymbol{s}) - \frac{\sigma}{1 + \boldsymbol{\gamma}^{+}\boldsymbol{s}}\boldsymbol{s} + \boldsymbol{\gamma}^{-}\boldsymbol{s}$$

Here $\boldsymbol{s}$ is a vector with each element standing for the abundance of one species on a single lattice point. The growth rate follows Gaussian distribution $\boldsymbol{\mu} \sim \mathcal{N}(0.6, 0.3^2)$. The interactions between two members followed Gaussian distribution $\boldsymbol{\gamma} \sim \mathcal{N}(0, 0.5^2)$. $\boldsymbol{\gamma}^{+}$ corresponds to the positive interactions in the $\boldsymbol{\gamma}$ matrix, and $\boldsymbol{\gamma}^{-}$ corresponds to the negative interactions (with negative values). The environmental stress for each member is set to $\sigma = 0.05$. We define the observables as the global abundances of the focal species, summed across the grid. ODE integration was performed for 50 a.u., and observable abundances were recorded every 1 a.u., yield time series with length of 50.

#### 1.3.3. Microbial communities with constant dispersal.

The gLV model with dispersal (**Fig. 3 b&d, S5b**) from a fixed species pool can generate highly complex dynamics when the community is large enough and interactions are strong enough[2]. We adopted the same model but further specified the growth rates for different species within the community:

$$\frac{d\boldsymbol{s}}{dt} = \boldsymbol{\mu s}(1 - \boldsymbol{s} - \boldsymbol{\gamma s}) + \boldsymbol{D}$$

The growth rate of each member follows a uniform distribution $\mu \sim U(0.5, 1.0)$. The interaction between two members follows a uniform distribution $\gamma \sim U(0, 0.8)$, with the positive values represent negative interactions. The dispersal for each member follows a uniform distribution $D \sim U(0, 2 \times 10^{-6})$.

In both “fixed” and “random” cases, the randomization of the initial abundances follows a uniform distribution of $s(0) \sim U(0, 0.2)$. ODE integration was performed for 300 a.u., and observable abundances were recorded every 6 a.u., yield time series with length of 50. To avoid observing trivial dynamics (e.g., exponential decay), we randomly chose the observables from the ones with final abundances > 0.05 at the end of one simulation instance.

#### 1.3.4. Microbial communities transferring plasmids.

To simulate complex microbial communities transferring plasmids (**Fig. 3 c&f, S5c**), we adopted a plasmid-centric model[3] to describe different microbial species $s_i$ and the abundance of species *i* carrying plasmid *j* $p_{ij}$.

$$\frac{ds_i}{dt} = \alpha_i \mu_i s_i c_i - D s_i$$

$$\frac{dp_{ij}}{dt} = \beta_{ij} \mu_i p_{ij} c_i + (s_i - p_{ij}) \sum_{k=1}^{m} \eta_{jki} p_{kj} - (\kappa_{ij} + D) p_{ij}$$

Here the effective burden on the total abundance of species *i* is described by $\alpha_i = s_i/(s_i + \sum_{j=1}^{n} p_{ij}\lambda_{ij})$, with $\lambda_{ij}$ being the burden of plasmid *j* on its host *i*. Effectively, if one assumes every type of plasmid is evenly distributed within a given species (no plasmid interactions like incompatibility), the growth burden of the subpopulation carrying plasmid *j* is penalized by carrying 100% of plasmid *j* and a fraction of other plasmids: $\beta_{ij} = s_i/(s_i(1 + \lambda_{ij}) + \sum_{k \neq j} p_{ik}\lambda_{ik})$. We adopted a niche-based model here by assuming there are 10 independent niches within the community, and different species within the same niche are competitively exclusive $c_i = e_i - \Sigma s_k$, where $e_i$ represents the carrying capacity of niche *i*. The plasmid *k* then conjugates between plasmid-free population of species *i* and plasmid-carrying populations of species *j* with rate $\eta_{jki}$, following a mass-action kinetics. Plasmid will also be lost at rate $\kappa_{ij}$. The entire community experiences constant dilution of $D$.

In our randomized simulations, the parameters all follow uniform distributions. The growth rate $\mu \sim U(0.2, 0.8)$, the plasmid burden $\lambda \sim U(1e-3, 3)$, the conjugation rate $\eta \sim U(1e-8, 0.02)$, and the niche capacities are drawn from $U(0, 1)$, and then normalized to have a summation of 1, the dilution rate is set to be 0.004. In our simulations, we fixed the initial abundance of every species $s \sim U(0.001, 1)$, and the background plasmids, but randomized the total abundance of the *n* observable plasmids in each simulation. We initialize the plasmid distribution by assigning $X \sim U(0, 1)$ as the maximum abundance of species *i* carrying *j*, and then assigning a scaling factor $Y \sim U(0, 1)$. The initial abundance of population $p_{ij}$ then becomes $p_{ij} = X \cdot Y$. The rationale for performing two rounds of randomization is to avoid the law of large numbers during one round of randomization, where the total abundance of plasmid *j* converges to a fixed value since it's averaged over 100 species. During the simulation, we took the plasmid abundance across the entire community as the target variable we are interested in. ODE integration was performed for 8400 a.u., and observable abundances were recorded every 168 a.u., yield time series with length of 50.

#### 1.4. Experimental data processing

OD600 and fluorescence intensities were measured for 192 cycles over 48 hours. Signals for the observables were obtained after subtracting the baseline measurements from the average of three blank controls per plate. Since the initial community size is very small, no significant signal was detected during the first 6 hours of incubation. So we focus on the target dynamics between 6 and 48 hours (168 time points). Cultures on the edges of the 384-wells were excluded from the dataset, as significant evaporation was observed. Cultures that did not grow (OD600 < 0.1) were also removed from the dataset. In addition, to ensure effective learning of the time series, we excluded three samples that either produced exceptionally high mEGFP or mCherry expression (maximum reading more than 3-fold higher than the rest of the samples). Altogether this yields a dataset of 887 datapoints.

For the sequenced endpoint community composition, we calculated the relative abundance of the 4 plasmid-carrying strains by the fraction of reads they take up in each sample.

#### 1.5. Literature data processing

##### 1.5.1. Long-term experimental microbial community dynamics

The first literature dataset we analyzed (**Fig. 5**) was from Fujita et al[4]. Fujita et al. inoculated a natural soil or pond water (water) microbiomes as the source microbiome into three types of media: A: oatmeal broth, B: oatmeal-peptone broth, or C: peptone broth, creating 6 distinct communities (Soil -A, -B, and -C, and Water -A, -B, and -C) with different dynamics, each with 8 replicates per community. 5-day pre-incubation was performed before the start of the experiment. The initial condition for each community varies across the 8 replicates. During the 110 days experiments, the communities were sampled every 24 hrs (taking 200 μL of sample for later sequencing analysis out of 1000 μL cultural media), and supplemented with 200 μL of fresh medium to each culture. 16S rRNA sequencing on Illumina platform was performed to quantify the change in the population size for each microbial ASV. During the bioinformatic analyses by the original study, samples with Pearson's coefficients of correlation between sequencing read numbers and standard DNA copy numbers less than 0.7 were removed. Samples with total reads less than 350 were also removed. The missing data are substituted by linear interpolation between known samples. During the experiment, 200 μL of fresh Medium B was accidently added to samples of Medium A on Day 82. So we only included the time series of Day 1 – 80 for Water-A and Soil-A communities. We included the entire 110-day time series for the rest 4 communities. We choose to analyze the dynamics of the relative abundance of the community.

**Data augmentation.** We first split the entire community dynamics time series (of 80 or 110 days) into 30-day segments, with each segment differing from one another by a minimum of 1 day. By doing so, we can generate 51 or 81 segments of the entire community dynamics. To create a larger dataset of *n* observables, we generated each datapoint in a rule-based manner as following: within a given segment (for example, day 1 - 30), we randomly selected one combination of *n* ASVs from the same replicate community. Since we are interested in learning the non-trivial dynamics, for each segment, we filtered out the ASVs whose time series $s_i(t)$ has cumulative abundance $\sum_{t=t_0}^{t_0+30} s_i(t) \leq 0.1$. For a given segment, if there are *N* ASVs with cumulative abundance > 0.1 within the same replicate community, the number of possible combinations is $\binom{N}{n}$. If $\binom{N}{n} \leq 300$, we exhausted all the combinations for this segment; otherwise, we randomly select 300 combinations for this segment, to keep the dataset balanced. For Water-C community, due to its low richness, the data augmentation results in small dataset, so we excluded this community for analysis.

**Quantification and forecast of community compositional shift.** Shift in community structure is quantified by the Bray-Curtis β-diversity between two compositions $\bar{\boldsymbol{s}}(t)$ and $\bar{\boldsymbol{s}}(t+1)$. To remove minor fluctuations, $\bar{\boldsymbol{s}}(t) = \frac{1}{5}\sum_{i=4}^{0} \boldsymbol{s}(t-i)$ and $\bar{\boldsymbol{s}}(t+1) = \frac{1}{5}\sum_{i=1}^{5} \boldsymbol{s}(t+i)$ are averaged across a 5-day window, before and after a given time point.

We tested the prediction accuracy of community compositional shift using different number of dominant ASVs (*n* = 1 – 10). Here we rank the ASVs by the number of reads they have across all the samples. For n = 1, only the time series of the most dominant ASV was encoded into the critical embedding to forecast the community shift.

#### 1.5.2. Human vaginal microbiome dynamics.

The second dataset we analyzed (**Fig. S18**) was the daily dynamics of 25 human vaginal microbiomes over 10 weeks, from Ravel et al.[5]. In this dataset, Ravel et al. measured the relative composition of vaginal microbiome samples from 25 subjects, among which 15 experienced symptomatic bacterial vaginosis (BV), 6 were diagnosed with asymptomatic BV, and 4 were healthy during the 10-week study. The relative composition of the sample communities was obtained by 16S rRNA amplicon sequencing via 454 pyrosequencing. Along with the vaginal swabs to sample the communities, pH value and menstruation are also documented daily, along with other meta data. We used linear interpolation between time points to fill in the missing samples across 70 days.

**Data augmentation.** We adopted the same strategy as in section 1.5.1. We first split the entire community dynamics time series (of 66 – 72 days) into 14-day segments, with each segment differing from one another by a minimum of 1 day. We generated each datapoint in the same rule-based manner: within a given segment, we randomly selected one combination of *n* taxa from the same subject. For each segment, we filtered out the taxa whose time series $s_i(t)$ has cumulative abundance $\sum_{t=t_0}^{t_0+14} s_i(t) \leq 0.1$. For a given segment, if there are *N* taxa with cumulative abundance > 0.1 within the same subject, the number of possible combinations is $\binom{N}{n}$. If $\binom{N}{n} \leq 100$, we exhausted all the combinations for this segment; otherwise, we randomly select 100 combinations for this segment, to keep the dataset balanced. We excluded the subject 49 for the low richness and small dataset size our method was able to generate.

### 2. Experimental methods

#### 2.1. Barcoded *E. coli* strains

62 *E. coli* strains were selected randomly from the Keio collection. The intergenic genomic site, referred to as Safe Site 3 (SS3) was selected as fixed locations for barcode insertions[6]. For plasmid construction, *E. coli* DH5α was used as cloning host. The pSS3gRNA plasmid that encodes the gRNA targeting the genomic locus of interest was constructed via recombination. Briefly, gRNA sequence with length of 20 bp was synthesized in primers and cloned into the pGRB backbone using the ClonExpress MultiS One-Step Cloning Kit (Vazyme, China). The pGRB plasmid was purchased from

Addgene (#71656). The assembled derivative plasmid pSS3gRNA was confirmed via Sanger sequencing.

We used CRISPR/Cas9-based genome editing system to insert the distinct barcode[7] for each Keio strain[8]. To achieve precise barcode insertions, we designed the donor DNAs with 400 bp homologous arms that flank the target site. To construct donor DNA, we first extracted the genome from an overnight culture of Keio strain. We next separately amplified two homologous arms from the genome DNA and the synthesized DNA fragment to be inserted (2 × Phanta max master mix, Vazyme, China), and then performed gel purification for the PCR products. We ligated three fragments by fusion PCR. The donor DNA and the pSS3gRNA plasmid expressing targeting gRNA were transformed into *E. coli* which harbored the pREDCas9 (Addgene, #71541) via electroporation. For preparing electrocompetent cells, we added 0.1 mM IPTG to the culture to induce the expression of λ-Red recombinase. Electrocompetent Keio strains were prepared using glycerol/mannitol density step centrifugation[9]. Bio-Rad MicroPulser was used for electroporation (0.2 cm cuvette, 2.50 kV, 25 μF, and 200 ohm). Cells after electroporation were immediately added into 500 μL SOC media and recovered for 1 h at 30°C. The recovered cells were plated on Lysogeny Broth (LB) agar plates containing 100 μg/mL spectinomycin and 100 μg/mL carbenicillin (Carb) at 30°C for 24 h. The correct transformant was validated by colony PCR and Sanger sequencing. For pSS3gRNA plasmid curing, the correct transformant was inoculated at 30°C for 12 h in the LB supplied with 2 g/L L-arabinose. Finally, the pREDCas9 in the strain was cured by cultivating at 42°C.

The Keio strains and corresponding barcodes are listed in Table S3.

#### 2.2. Community assembly experiment leveraging limiting dilution

We included the 62 barcoded Keio strains in the community assembly process. Among the 62 Keio strains, 4 strains were randomly selected as plasmid carriers. The plasmids each has PBR322 replication origin, ampicillin resistance, and fluorescence marker under pBAD promoter (Addgene, mEGFP: #54622, mCherry: #54630, mTagBFP2: #34632; LSSmOrange: #37129). The four strains (K1 – K4) were made chemically competent using calcium chloride and transformed via heat shock at 42 °C for 1 minute, followed by recovery in SOC medium.

Individual strains were cultured in LB directly from frozen glycerol stock for 18 hours before community assembly. The plasmid-carrying strains were cultured in LB supplemented with 100 ug/mL Carb and 0.2% w/v L-arabinose. The plasmid-free strains were cultured in LB supplemented with 50 ug/mL Kanamycin (Kan).

Equal volumes of the 58 plasmid-free Keio strains and 4× volumes of the 4 plasmid-carrying strains were mixed. The mixture was further diluted into LB supplemented with 2 ug/mL Carb and 0.2 % w/v L-arabinose. Three independent plates were cultured on different days, two of which used the dilution ratios of $4\times10^8$ for plasmid-free strains and ~ $1\times10^8$ for plasmid-carrying strains from the original overnight culture, and one used $2\times10^8$ for plasmid-free strains and ~ $5\times10^7$ for plasmid-carrying strains.

100 uL of the above culture was dispensed into each well of Thermo Scientific Nunc 384 transparent microplate. The cells for each strain per well follow Poisson distribution, resulting in random assembly of communities. To reduce evaporation, the plate was loaded with the lid into Tecan Infinity microplate reader. The cultures were incubated at 37 ℃. OD600 and fluorescence intensities are measured 15 minutes with periodical shaking (orbital, 5 s, 2 mm) for 48 hours. The excitation and emission frequencies for each fluorescent protein were chosen as follow: mCherry: excitation 561 nm, emission

600 nm; mEGFP: excitation 488 nm, emission 516 nm; mTagBFP2: excitation 399 nm, emission 454nm; LSSmOrange: excitation 437 nm, emission 572 nm.

To avoid edge effect, we excluded the samples on the edges of the 384-well plates during data processing.

#### 2.3. Next-generation sequencing (NGS) for endpoint community composition

After incubation of 48 hours, the cultures were stored in 384 well plates at -80 ℃. Upon the preparation of NGS library, the communities were boiled at 98 ℃ for 20 minutes. The barcodes within each community ere amplified through PCR using DNA polymerase (2 × Phanta max master mix, Vazyme, China). The primers for PCR each contain unique 8 bp index for multiplexing the samples for sequencing. Products from PCR were pooled together and run on 2% agarose gel. DNA was extracted and purified from the gel using the Zymoclean Gel DNA Recovery kit, based on the manufacturer's instructions. The cleaned PCR product was compatible with standard Illumina sequencing platforms. The DNA libraries were denatured, diluted, and mixed with an equal amount of Phi-X spike-in based on standard Illumina protocols for preparation of 16S libraries on the Illumina Miniseq. The final libraries contained 50% of Phi-X to ensure library diversity. The libraries were sequenced using 151-base pair paired-end reads on Illumina MiniSeq. The final library from PCR is:

5'-AATGATACGGCGACCACCGAGATCTACACXXXXXXXXAACACTCTTTCCC
TACACGACGCTCTTCCGATCTNNNNNNNNNNNNNNNNNNNNNNNNNNNNAGATCG
GAAGAGCACACGTCTGAACTCCAGTCACXXXXXXXXTCTCGTATGCCGTCTTCTGC
TTG-3'

Here, N's represent barcode sequences, and X's represent sample indexes.

#### 2.4. Sequencing data analysis

A customized Galaxy workflow was used to demultiplex the samples based on the index sequences from the paired-end reads and to count the barcodes of each community. The following tools were used: Trim sequences (Galaxy Version 1.0.2+galaxy0), Barcode Splitter (Galaxy Version 1.0.1), FASTQ joiner (Galaxy Version 2.0.1.1+galaxy0).

### 2.5. Estimation of average cell number in a community

We estimated the average initial cell number of each strain in each community based on Poisson distribution and the endpoint NGS result. We considered yielding 0 reads from the NGS result as the strain being absent. Assuming uniform dispensing across the entire 384 well plate, we calculated the fraction of wells that's missing a particular strain, averaged across the 58 background strains or the 4 target strains. The averaged missing fraction $P_0$, can be used calculated the Poisson parameter $\lambda = -logP_0$. Both including and excluding the edge wells yielded the same estimate.

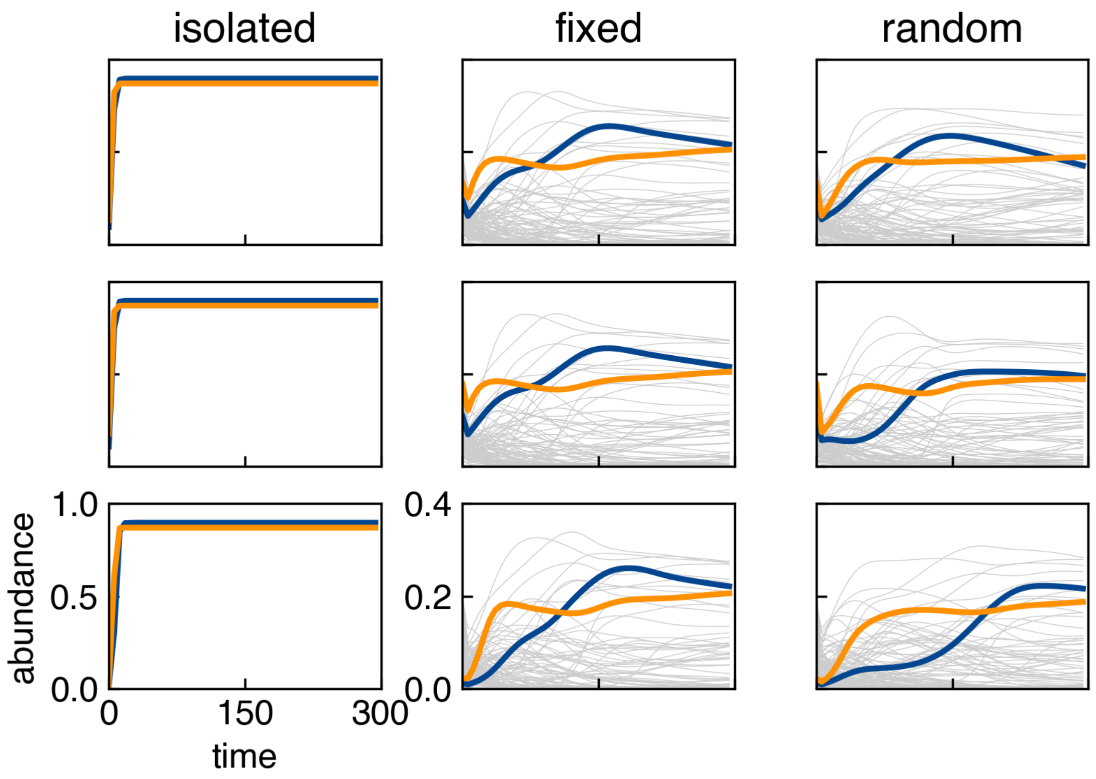


**Figure S1. Dynamics of the 2 observables (blue and orange) and the 98 background populations (grey) under different simulation conditions.**

Simulations are performed when the observables are 1) isolated from the background populations; 2) co-cultured with a background community with fixed initial conditions across different simulations; and 3) co-cultured with a background community with randomized initial conditions across different simulations.

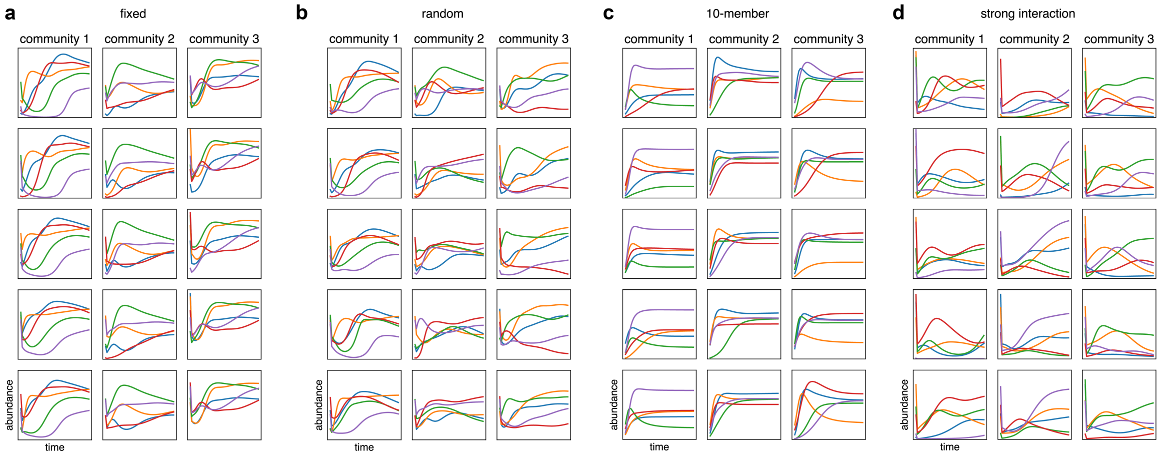


**Figure S2. Example time series of *n* = 5 observables generated by the bounded gLV model.**

a) 100-member communities in “fixed” case simulations.

b) 100-member communities in “random” case simulations.

c) 10-member communities in “random” case simulations.

d) 100-member communities with stronger interactions in “random” case simulations.

For each simulation case, 3 randomized communities are generated, and 5 examples are shown for each column per community.

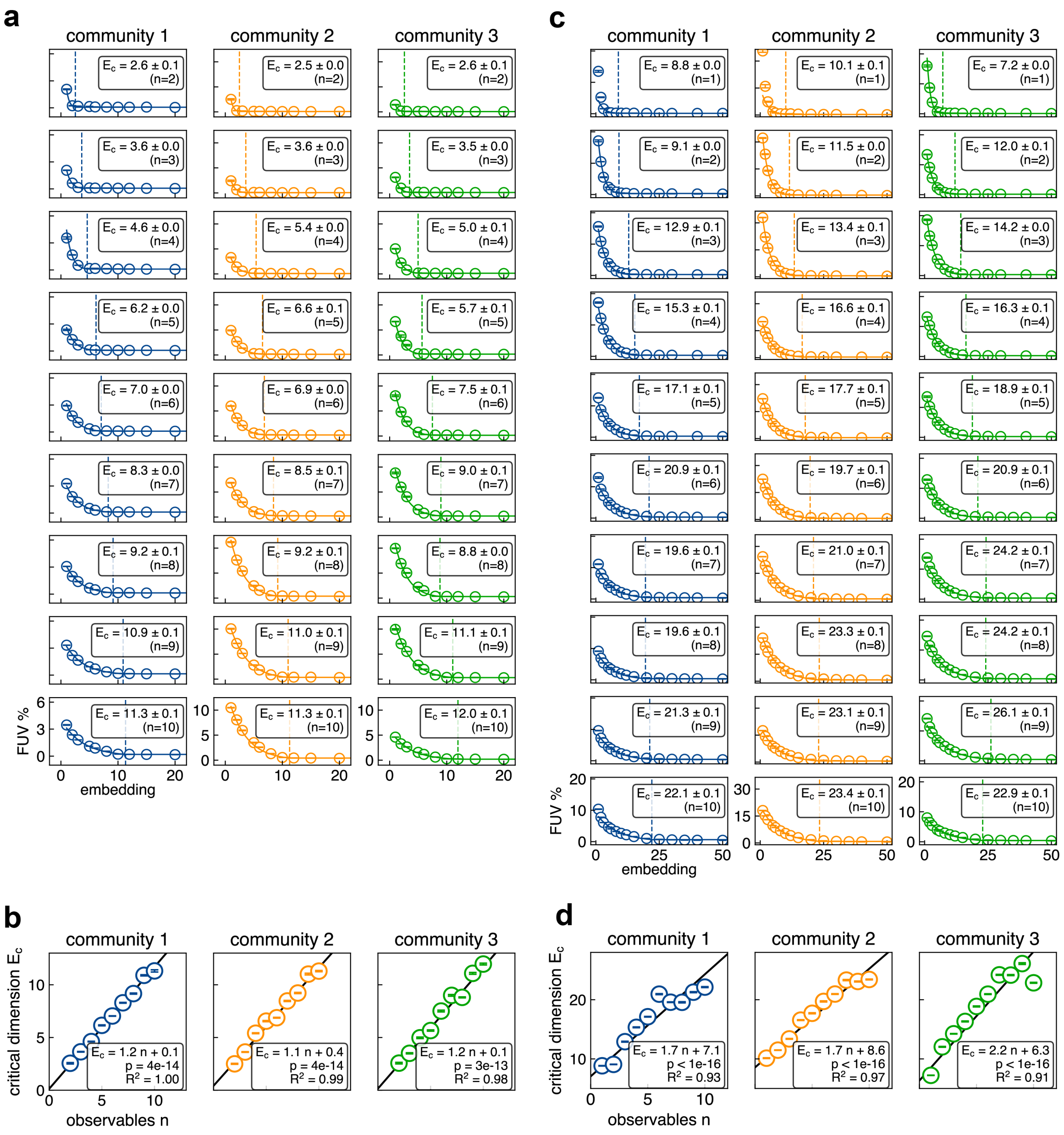


**Figure S3. Quantification of the critical dimension of the observables for the bounded gLV model.**

a) The FUV (mean ± SE, 5 replicates) between simulated and reconstructed time series data in the "fixed" case by VAEs with different embedding dimensions. σ-weighted regression was performed to fit the datapoints to a threshold-exponential function to estimate the critical dimension *Ec* (mean ± SE).

b) $E_c$ (mean ± SE) linearly correlates with *n* for the “fixed” case (σ-weighted least-squares regression. Community 1: slope 1.157 ± 0.006 SE, $p = 4\times10^{-14}$; weighted $R^2 = 0.996$. Community 2: slope 1.120 ± 0.006 SE, $p = 4\times10^{-14}$; weighted $R^2 = 0.986$. Community 3: slope 1.151 ± 0.008 SE, $p = 3\times10^{-13}$; weighted $R^2 = 0.979$.)

c) The FUV (mean ± SE, 5 replicates) between simulated and reconstructed time series data in the “random” case by VAEs with different embedding dimensions. σ-weighted regression was performed to fit the datapoints to a threshold-exponential function to estimate the critical dimension *E*c (mean ± SE).

d) $E_c$ (mean ± SE) linearly correlates with *n* for the “random” case (σ-weighted least-squares regression. Community 1: slope 1.725 ± 0.006 SE, $p < 1\times10^{-16}$; weighted $R^2 = 0.925$. Community 2: slope 1.667 ± 0.006 SE, $p < 1\times10^{-16}$; weighted $R^2 = 0.971$. Community 3: slope 2.180 ± 0.006 SE, $p < 1\times10^{-16}$; weighted $R^2 = 0.912$.)

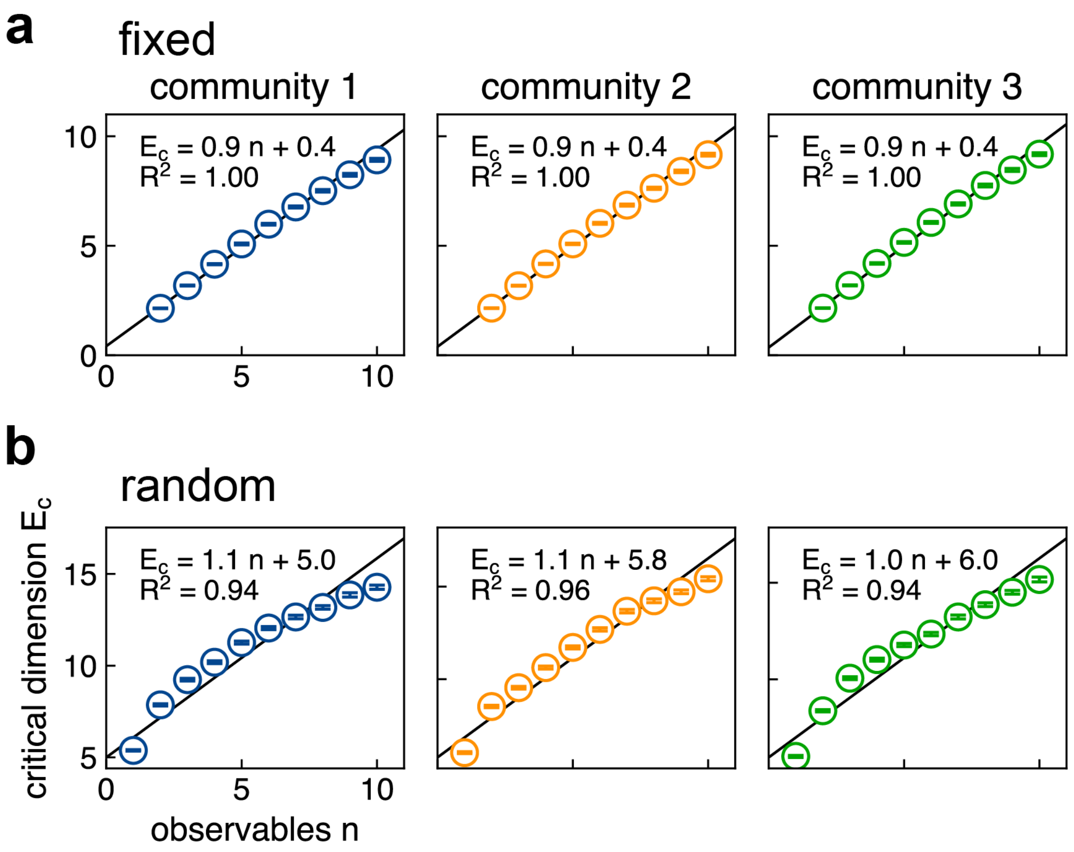


**Figure S4. Maximum likelihood estimation confirms the linear relationship between the intrinsic dimension and the number of observables.**

a) $E_c$ (mean ± bootstrap averaged standard deviation) linearly correlates with *n* for the "fixed" case (σ-weighted least-squares regression. Community 1: slope 0.898 ± 0.005 SE, $p = 2\times10^{-14}$; weighted $R^2 = 0.995$. Community 2: slope 0.913 ± 0.004 SE, $p = 1\times10^{-14}$; weighted $R^2 = 0.997$. Community 3: slope 0.927 ± 0.005 SE, $p = 2\times10^{-14}$; weighted $R^2 = 0.997$.)

b) $E_c$ (mean ± bootstrap averaged standard deviation) linearly correlates with *n* for the "random" case (σ-weighted least-squares regression. Community 1: slope 1.085 ± 0.008 SE, $p = 1\times10^{-14}$; weighted $R^2 = 0.936$. Community 2: slope 1.075 ± 0.008 SE, $p = 2\times10^{-14}$; weighted $R^2 = 0.956$. Community 3: slope 1.030 ± 0.008 SE, $p = 2\times10^{-14}$; weighted $R^2 = 0.936$.)

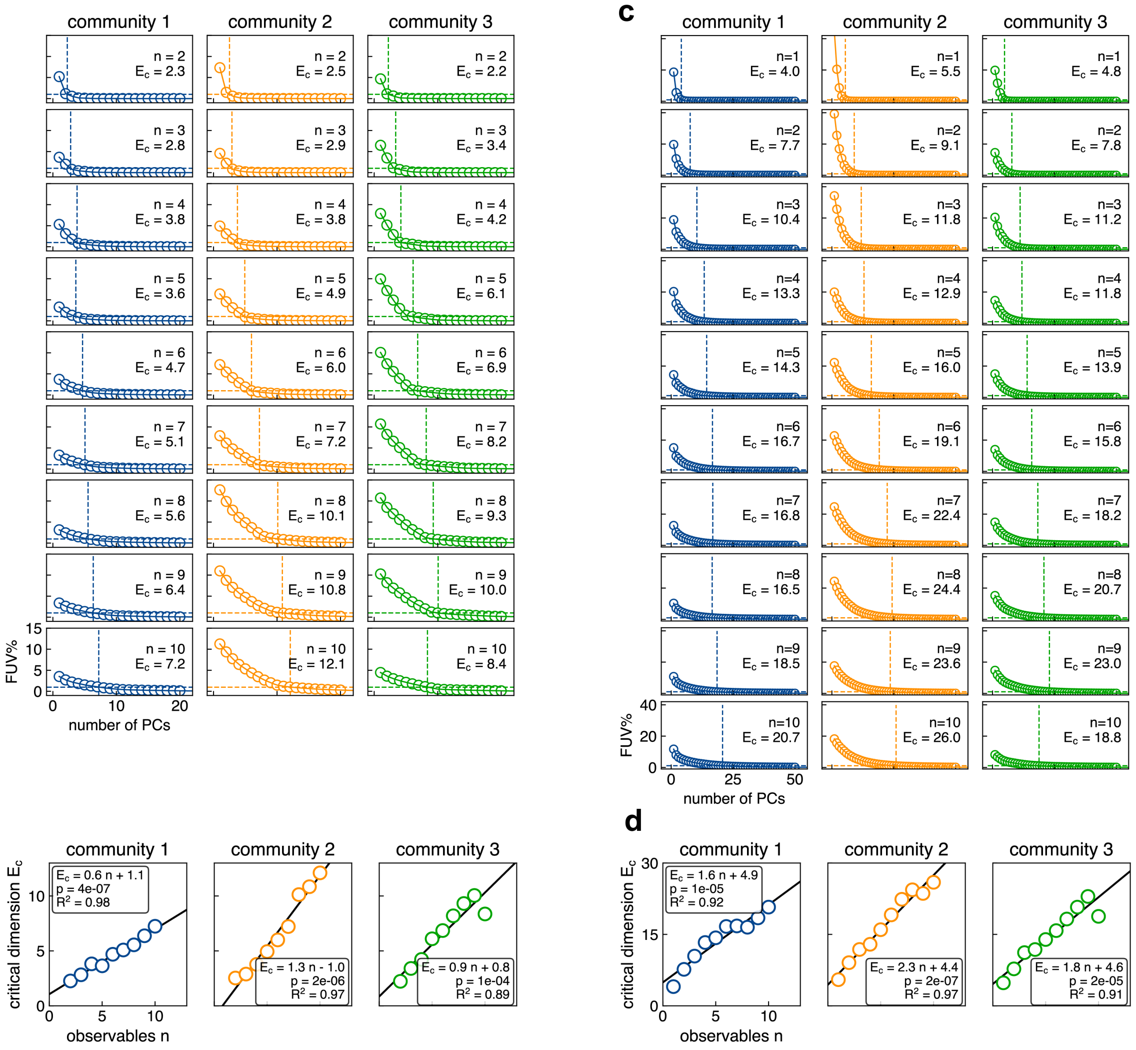


**Figure S5. Principal component analysis (PCA) identified similar linear scalings between $E_c$ and $n$ for the bounded gLV model.**

a) Observable time series in the “fixed” case are flattened and decomposed using PCA with varying numbers of principal components. Each reduced representation is reconstructed, and the FUV is computed between the reconstructed and original time series. Critical dimension $E_c$ is defined as the number of principle components with FUV = 1% based on linear interpolation.

b) $E_c$ linearly correlates with $n$ for the “fixed” case (Community 1: slope 0.591 ± 0.033 SE, $p = 4\times10^{-7}$; weighted $R^2$ = 0.979. Community 2: slope 1.286 ± 0.088 SE, $p$ =

$2\times10^{-6}$; weighted $R^2$ = 0.986. Community 3: slope 0.946 ± 0.124 SE, $p = 1\times10^{-4}$; weighted $R^2$ = 0.892.)

c) FUV between the reconstructed and original time series in the “random” case. Critical dimension $E_c$ is defined as the number of principle components with FUV = 1% based on linear interpolation.

d) $E_c$ linearly correlates with $n$ for the “random” case (Community 1: slope 1.632 ± 0.173 SE, $p = 1\times10^{-5}$; weighted $R^2$ = 0.918. Community 2: slope 2.304 ± 0.142 SE, $p = 2\times10^{-7}$; weighted $R^2$ = 0.986. Community 3: slope 1.825 ± 0.204 SE, $p = 2\times10^{-5}$; weighted $R^2$ = 0.909.)

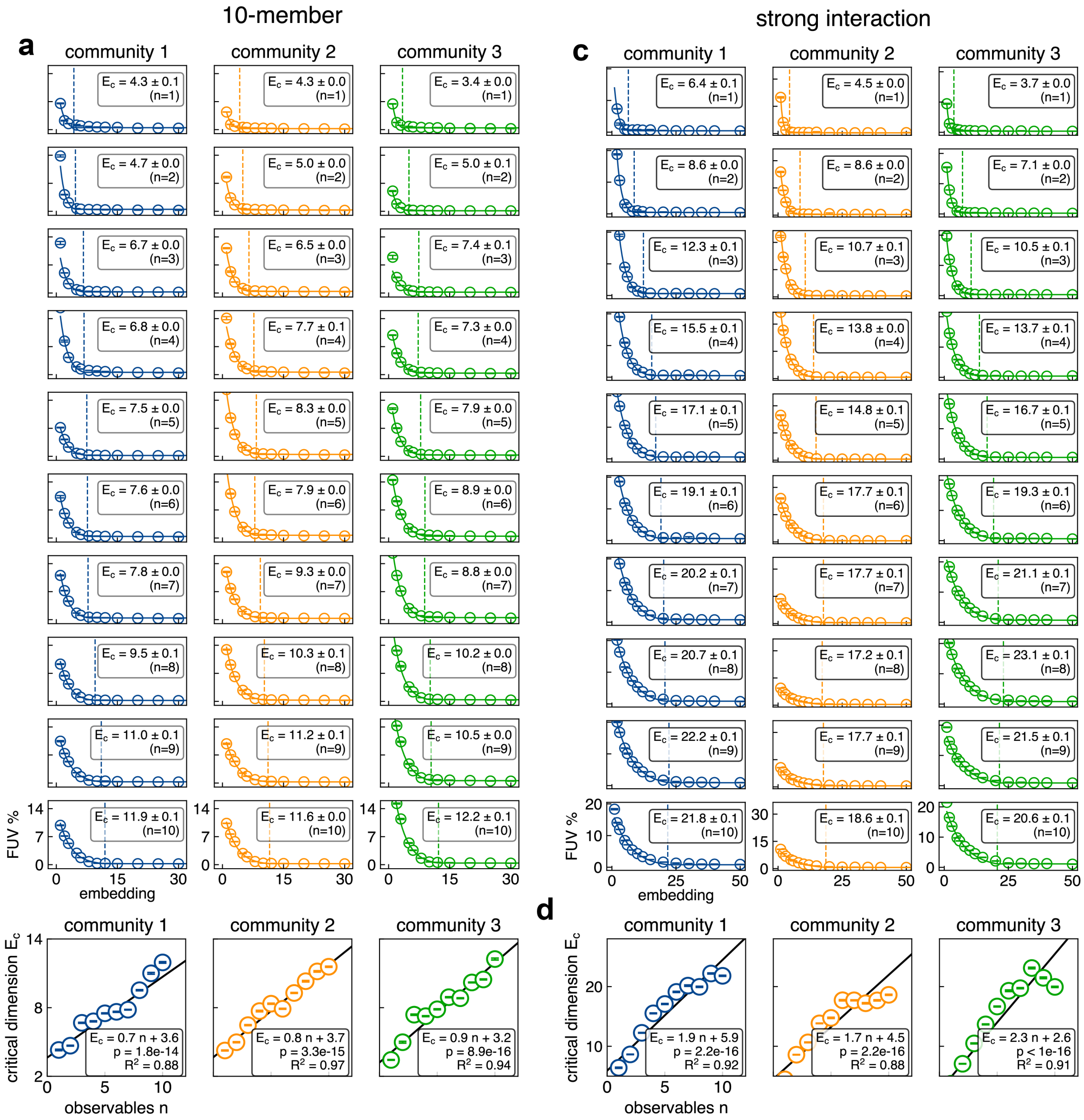


**Figure S6. Linear correlation between $E_c$ and n consistently holds regardless of community size and interaction strength.**

a) The FUV (mean ± SE, 5 replicates) between simulated and reconstructed time series data in the “random” case in 10-member communities by VAEs with different embedding dimensions. σ-weighted regression was performed to fit the datapoints to a threshold-exponential function to estimate the critical dimension *E*c (mean ± SE).

b) $E_c$ (mean ± SE) linearly correlates with $n$ (σ-weighted least-squares regression. Community 1: slope 0.706 ± 0.006 SE, $p = 2\times10^{-14}$; weighted $R^2 = 0.883$. Community 2: slope 0.807 ± 0.005 SE, $p = 3\times10^{-15}$; weighted $R^2 = 0.968$. Community 3: slope 0.871 ± 0.005 SE, $p = 9\times10^{-16}$; weighted $R^2 = 0.944$).

c) The FUV (mean ± SE, 5 replicates) between simulated and reconstructed time series data in the “random” case in 100-member communities with stronger interactions by VAEs with different embedding dimensions. σ-weighted regression was performed to fit the datapoints to a threshold-exponential function to estimate the critical dimension $Ec$ (mean ± SE).

d) $E_c$ (mean ± SE) linearly correlates with $n$ (σ-weighted least-squares regression. Community 1: slope 1.855 ± 0.008 SE, $p = 2\times10^{-16}$; weighted $R^2 = 0.923$. Community 2: slope 1.744 ± 0.007 SE, $p = 2\times10^{-16}$; weighted $R^2 = 0.875$. Community 3: slope 2.290 ± 0.008 SE, $p < 1\times10^{-16}$; weighted $R^2 = 0.915$).

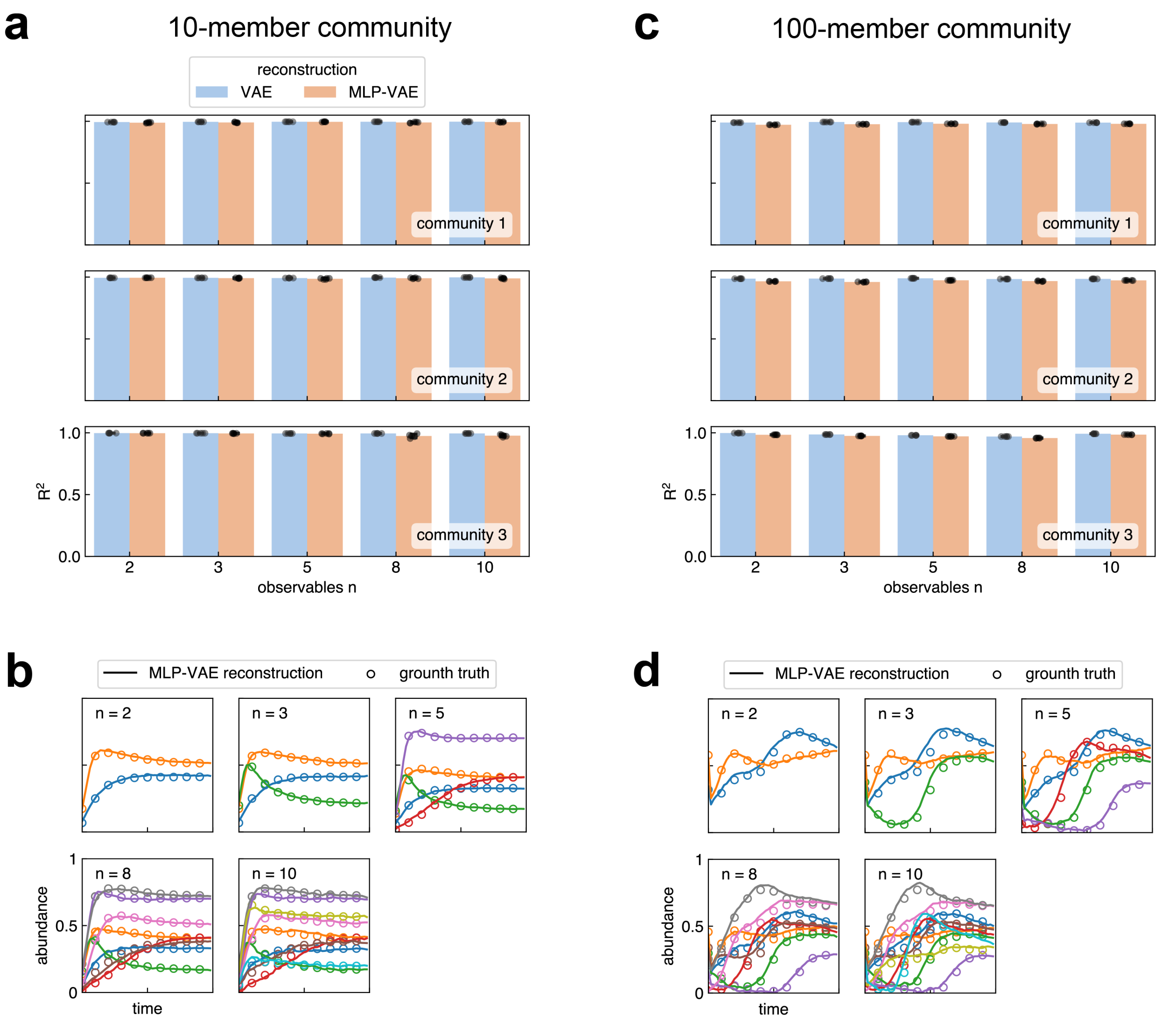


**Figure S7. MLP-VAE models can map the initial abundances of the communities to the observable dynamics.**

a) The MLP model maps the initial abundances of the 10-member community to the critical embeddings, which were further decoded into the observable dynamics by the VAE model. The MLP-VAE models achieve comparable performance (community 1: $R^2$ = 0.992 ± 0.004 SD; community 2: $R^2$ = 0.990 ± 0.004 SD; community 3: $R^2$ = 0.987 ± 0.012 SD. 5 different $n$ × 5 training replicates) in target dynamics prediction as the VAE reconstruction (community 1: $R^2$ = 0.996 ± 0.001 SD; community 2: $R^2$ = 0.994 ± 0.002 SD; community 3: $R^2$ = 0.996 ± 0.001 SD. 5 different $n$ × 5 training replicates) .

b) Examples of ground truth (open circles) and MLP-VAE predictions (curve) of n observables in a 10-member community (Community 1).

c) The MLP model maps the initial abundances of the 100-member community to the critical embeddings, which were further decoded into the observable dynamics by the VAE model. The MLP-VAE models achieve comparable performance (community 1: $R^2$ = 0.977 ± 0.003 SD; community 2: $R^2$ = 0.968 ± 0.005 SD; community 3: $R^2$ = 0.974 ± 0.010 SD. 5 different *n* × 5 training replicates) in target dynamics prediction as the VAE reconstruction (community 1: $R^2$ = 0.991 ± 0.002 SD; community 2: $R^2$ = 0.987 ± 0.005 SD; community 3: $R^2$ = 0.985 ± 0.010 SD. 5 different *n* × 5 training replicates) .

d) Examples of ground truth (open circles) and MLP-VAE predictions (curve) of n observables in a 100-member community (Community 1).

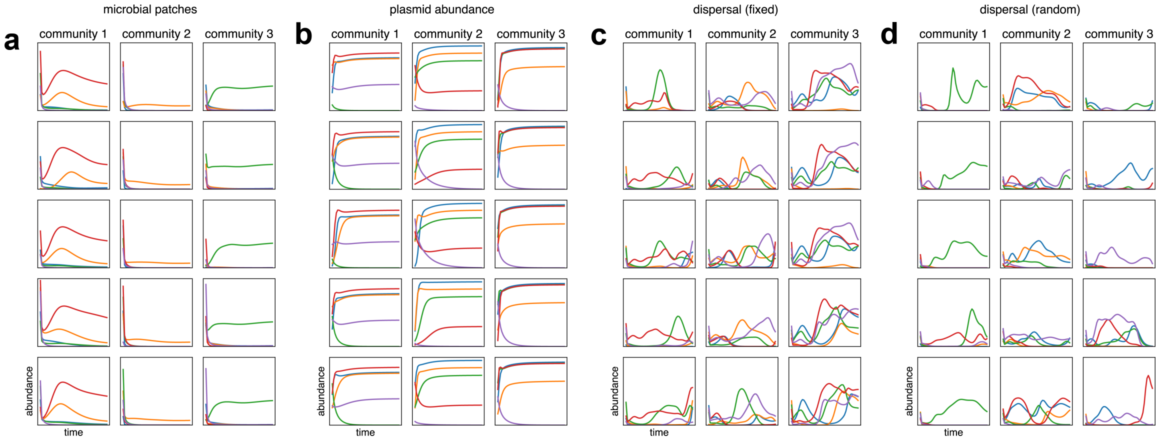


**Figure S8. Example time series of $n$ = 5 observables in different microbial ecosystems.**

a) Microbial patchy dynamics.

b) Plasmid dynamics.

c) Communities with constant dispersal in “fixed” case simulations.

d) Communities with constant dispersal in “random” case simulations.

For each simulation case, 3 randomized communities are generated, and 5 examples are shown for each column per community.

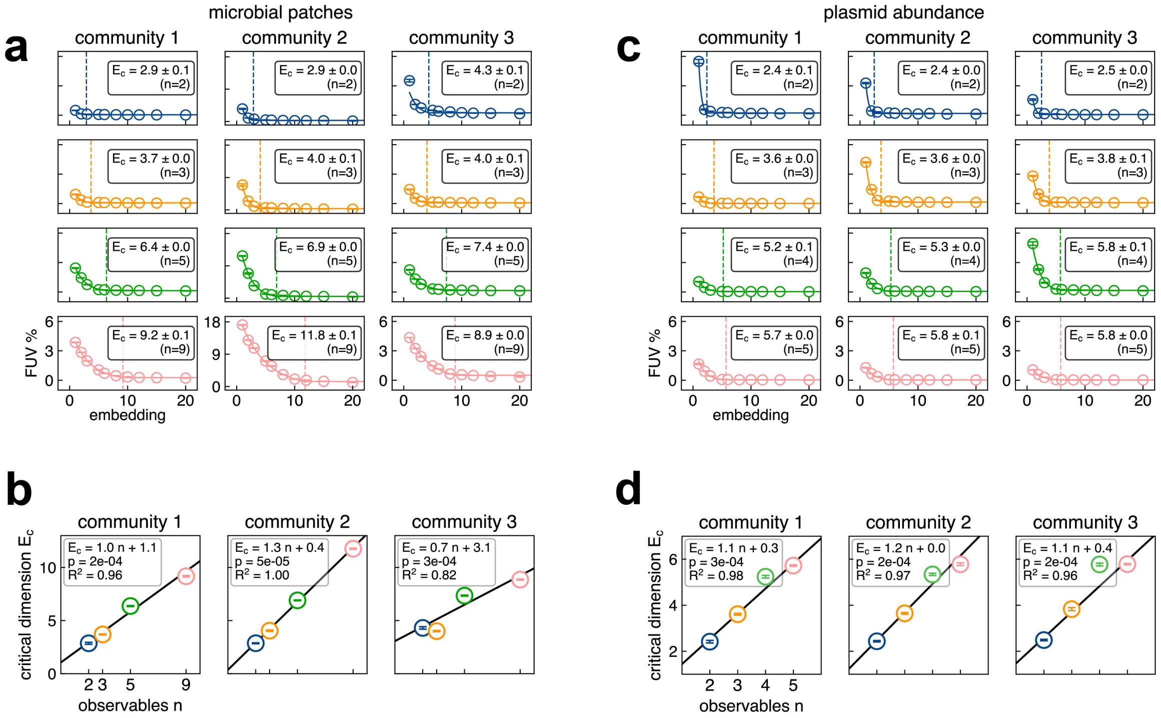


**Figure S9. Quantification of the critical dimension of the observables in microbial patchy dynamics and plasmid dynamics.**

a) The FUV (mean ± SE, 5 replicates) between simulated and reconstructed observable patchy dynamics by VAEs with different embedding dimensions. σ-weighted regression was performed to fit the datapoints to a threshold-exponential function to estimate the critical dimension *E*c (mean ± SE).

b) $E_c$ (mean ± SE) linearly correlates with *n* for microbial patchy dynamics (σ-weighted least-squares regression. Community 1: slope 0.955 ± 0.009 SE, $p = 2 \times 10^{-4}$; weighted $R^2 = 0.963$. Community 2: slope 1.280 ± 0.006 SE, $p = 5 \times 10^{-5}$; weighted $R^2 = 0.999$. Community 3: slope 0.685 ± 0.011 SE, $p = 3 \times 10^{-4}$; weighted $R^2 = 0.822$.)

c) The FUV (mean ± SE, 5 replicates) between simulated and reconstructed plasmid dynamics. σ-weighted regression was performed to fit the datapoints to a threshold-exponential function to estimate the critical dimension *E*c (mean ± SE).

d) $E_c$ (mean ± SE) linearly correlates with *n* for plasmid dynamics (σ-weighted least-squares regression. Community 1: slope 1.094 ± 0.018 SE, $p = 3\times10^{-4}$; weighted $R^2 = 0.976$. Community 2: slope 1.228 ± 0.019 SE, $p = 2\times10^{-4}$; weighted $R^2 = 0.972$. Community 3: slope 1.119 ± 0.014 SE, $p = 2\times10^{-4}$; weighted $R^2 = 0.955$.)

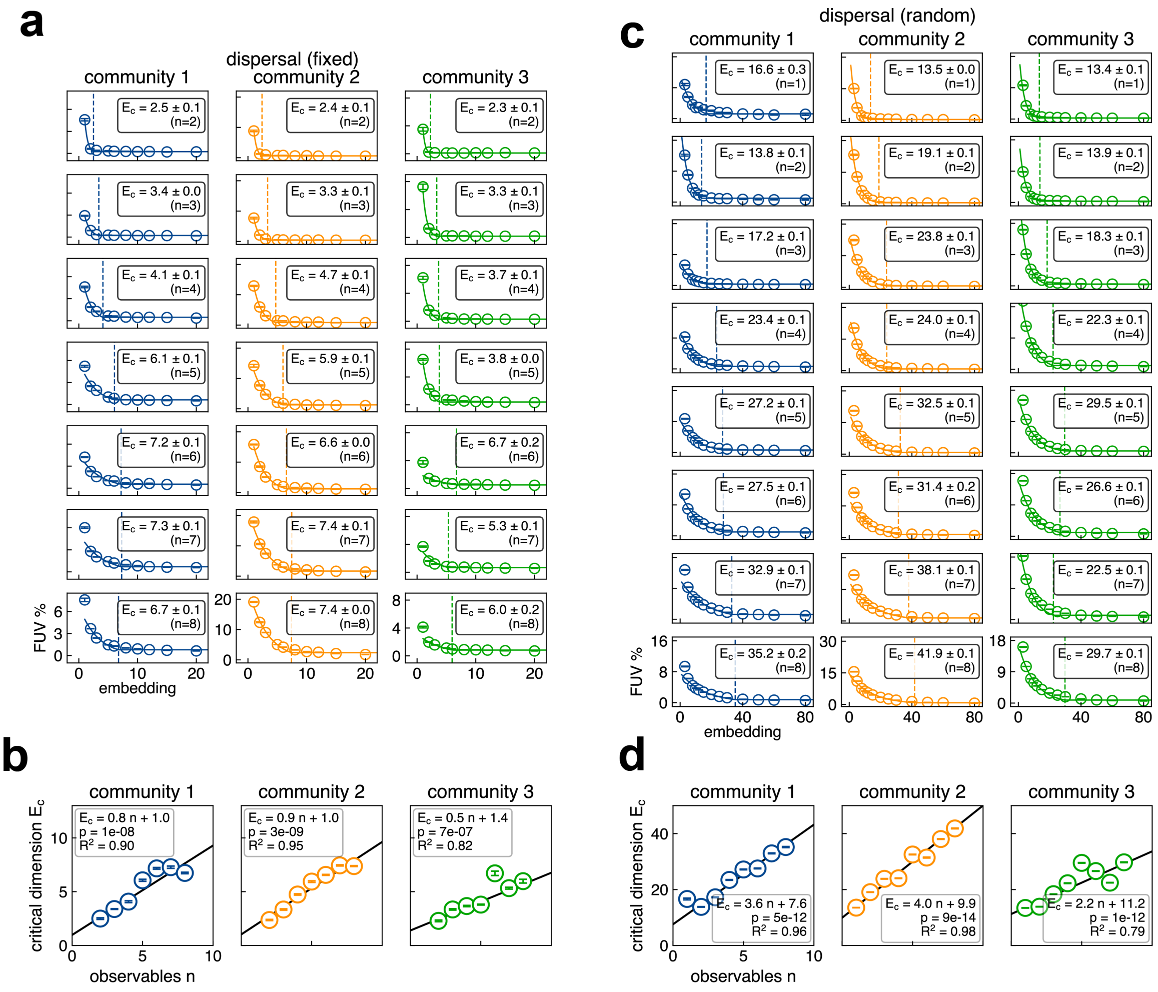


**Figure S10. Quantification of the critical dimension of the observables in community dynamics with continuous immigration.**

a) The FUV (mean ± SE, 5 replicates) between simulated and reconstructed observable dynamics in the "fixed" case of the gLV simulation with constant dispersal. σ-weighted regression was performed to fit the datapoints to a threshold-exponential function to estimate the critical dimension *E*c (mean ± SE).

b) $E_c$ (mean ± SE) linearly correlates with *n* (σ-weighted least-squares regression. Community 1: slope 0.832 ± 0.012 SE, $p = 1\times10^{-8}$; weighted $R^2$ = 0.897. Community

2: slope 0.861 ± 0.009 SE, $p = 3\times10^{-9}$; weighted $R^2 = 0.948$. Community 3: slope 0.537 ± 0.018 SE, $p = 7\times10^{-7}$; weighted $R^2 = 0.819$.)

c) The FUV (mean ± SE, 5 replicates) between simulated and reconstructed observable dynamics in the “fixed” case of the gLV simulation with constant dispersal. σ-weighted regression was performed to fit the datapoints to a threshold-exponential function to estimate the critical dimension *E*c (mean ± SE).

d) $E_c$ (mean ± SE) linearly correlates with *n* (σ-weighted least-squares regression. Community 1: slope 3.568 ± 0.023 SE, $p = 5\times10^{-12}$; weighted $R^2 = 0.958$. Community 2: slope 4.017 ± 0.013 SE, $p = 9\times10^{-14}$; weighted $R^2 = 0.978$. Community 3: slope 2.243 ± 0.011 SE, $p = 1\times10^{-12}$; weighted $R^2 = 0.790$.)

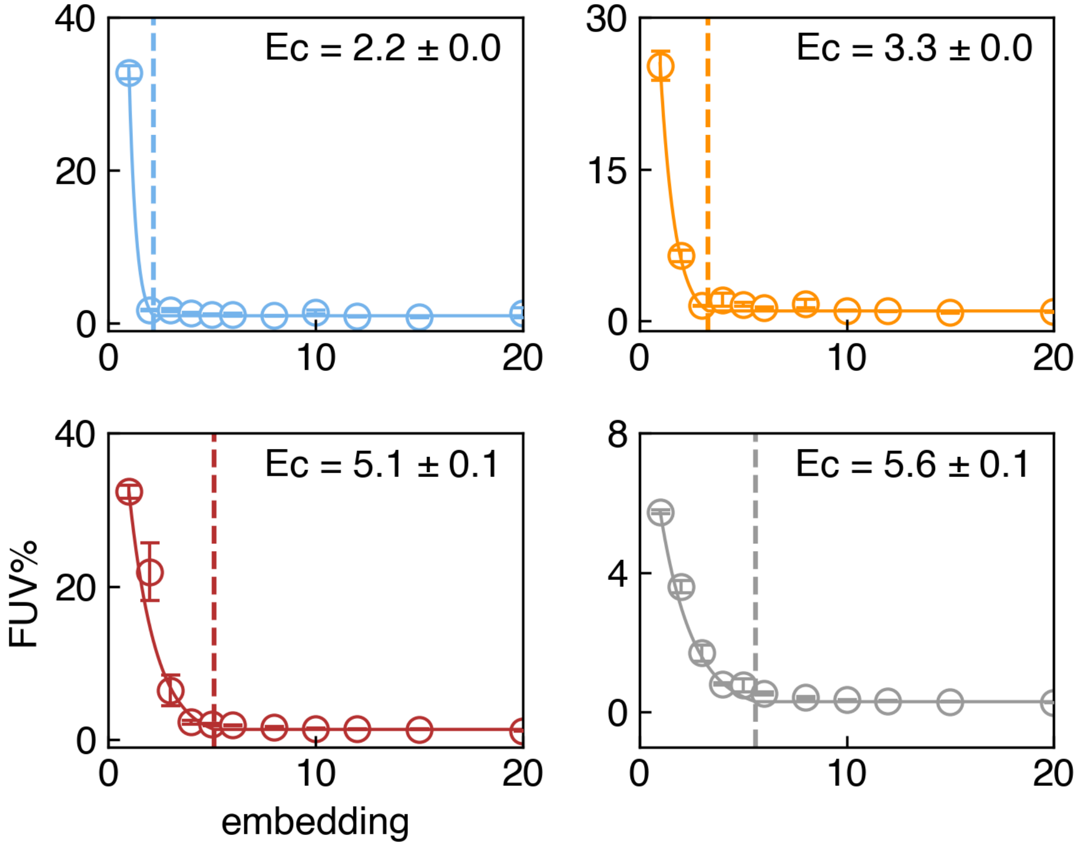


**Figure S11. Quantification of the critical dimension of the observables during community assembly.**

The FUV (mean ± SE, 5 training replicates) between experimental and reconstructed time series data by VAEs with different embedding dimensions. σ-weighted regression was performed to fit the datapoints to a threshold-exponential function to extract the critical dimension *E*c (mean ± SE). Here *n* = 2 corresponds to mEGFP and mTagBFP2, *n* = 3 includes LSSmOrange, *n* = 4 includes mCherry, and *n* = 5 includes OD.

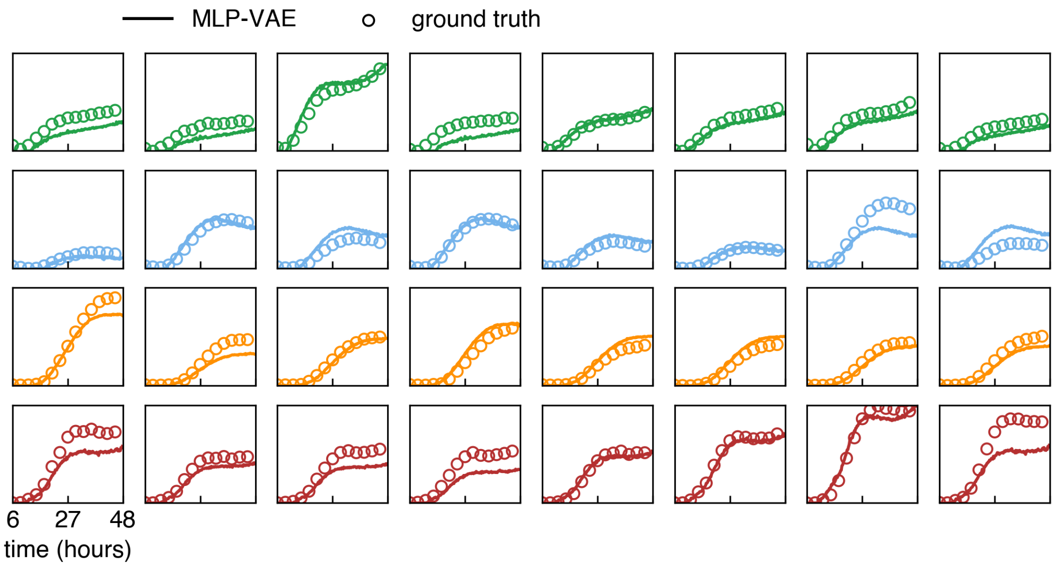


**Figure S12. Fluorescence dynamics can be inferred based on the endpoint abundances of plasmid-carrying strains by the MLP-VAE model.**

Example time series of inferred (curve) and ground truth (open circles) fluorescence dynamics by the MLP-VAE model. Row 1: mEGFP, Row 2: mTagBFP2, Row 3: LSSmOrange, Row 4: mCherry.

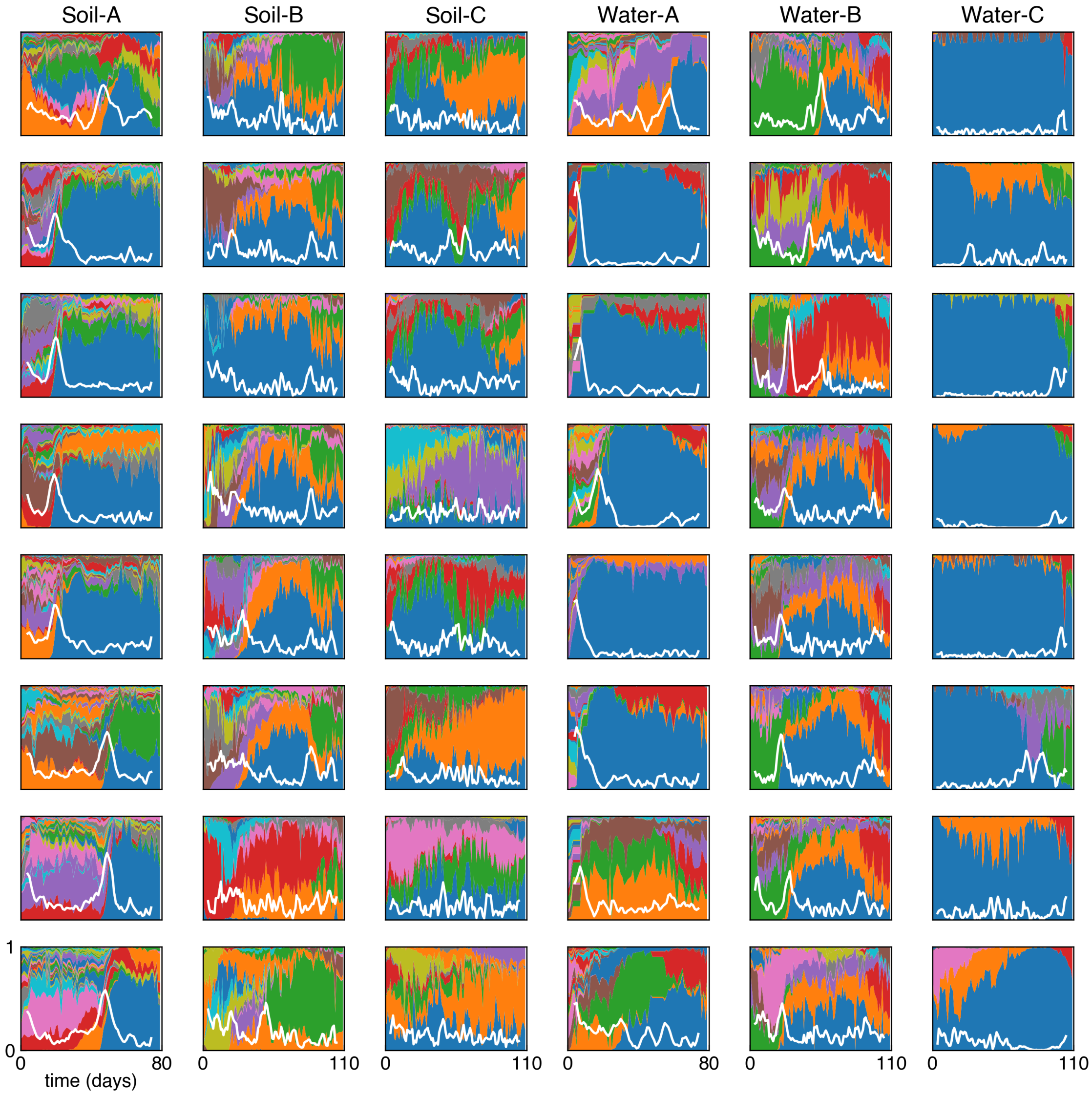


**Figure S13. Community dynamics of environment-derived microbiomes.**

Composition of 8 replicate communities over 80 days (for Soil-A and Water-A communities) and over 110 days (for Soil -B, -C, Water -B, -C communities). The white curves show the calculated Bray-Curtis β-diversity over time.

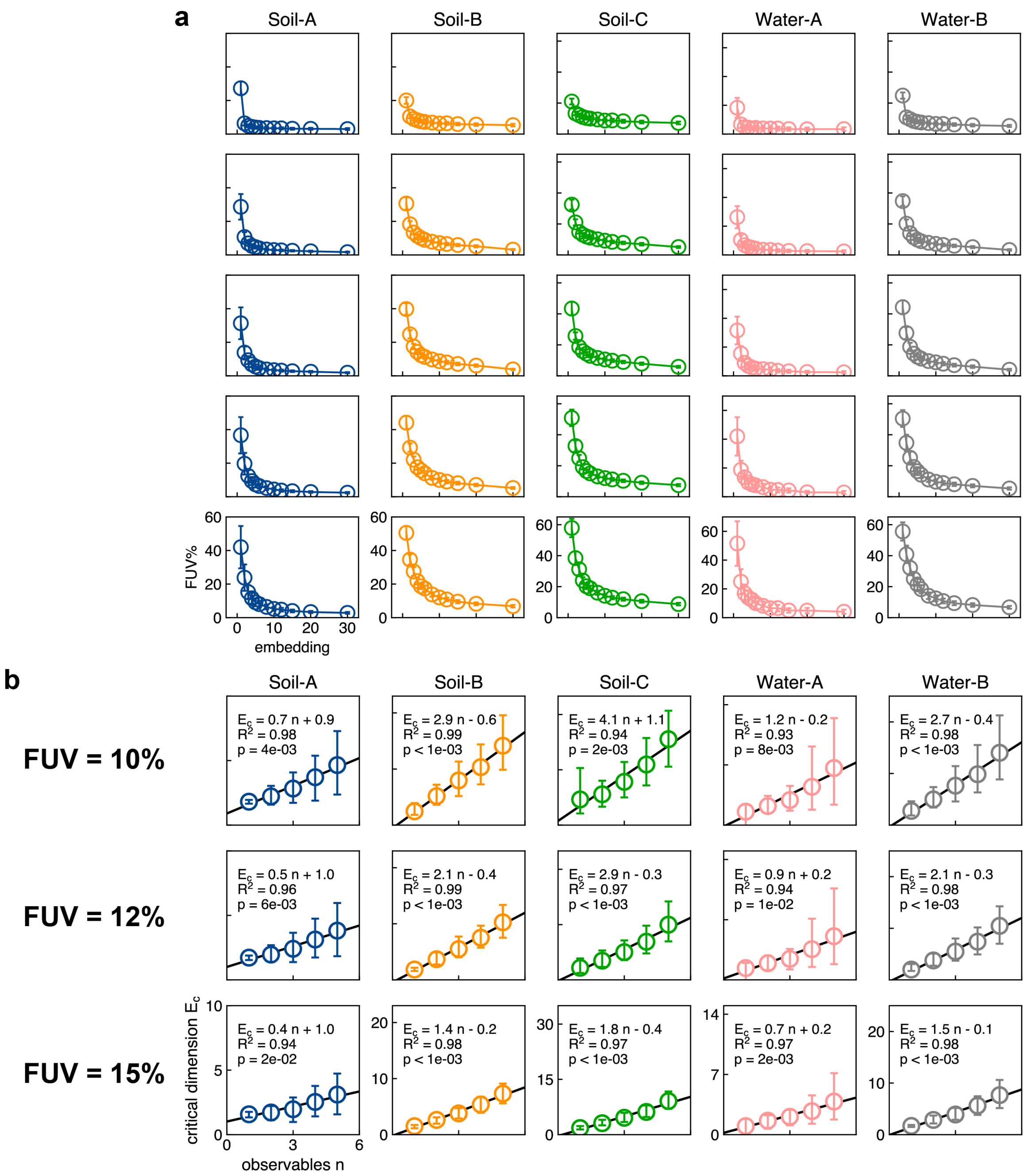


**Figure S14. Time series of experimental microbial dynamics can be represented by low-dimensional embeddings.**

a) Dependence of FUV (3 training replicates are averaged for each cross-validation fold, mean ± SE across 8-fold cross-validation was shown) on the embedding dimension.

b) Linear correlations between $E_c$ and $n$ consistently holds for all five communities for FUV thresholds of 10%, 12%, and 15%. Each point represents the mean $E_c$ from bootstrapping interpolation (1,000 times), together with 95% CI. Black line indicates the best linear fit to mean $E_c$ across $n$, with reported $R^2$ and bootstrapped p-value testing the slope is significantly greater than zero.

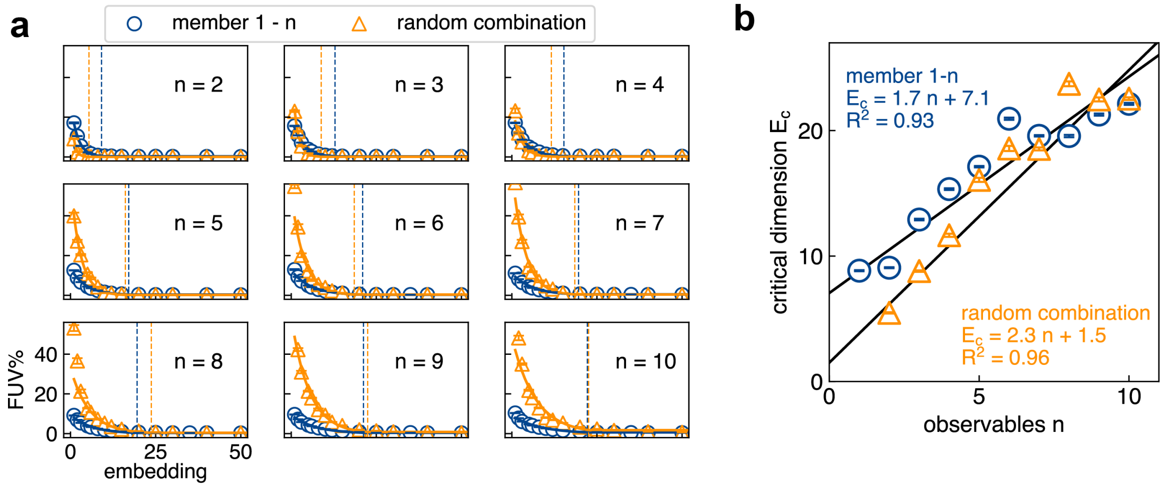


**Figure S15. The critical dimension of datasets generated from random combination of simulated observable dynamics.**

a) The FUV (mean ± SE, 5 replicates) between ground truth and reconstruction for the "random case" (blue, same as Fig.2d) and for the random-combination-augmented datasets (orange). σ-weighted regression was performed to fit the datapoints to a threshold-exponential function to estimate the critical dimension *E*c (mean ± SE). $E_c$ is indicated by the dashed lines.

b) $E_c$ (mean ± SE) linearly correlates with *n* for both cases (σ-weighted least-squares regression. Member 1 – *n*, same as Fig. 2d: slope 1.725 ± 0.006 SE, $p < 1\times10^{-16}$; weighted $R^2 = 0.925$. Random combination: slope 2.332 ± 0.012 SE, $p = 2\times10^{-14}$; weighted $R^2 = 0.960$.)

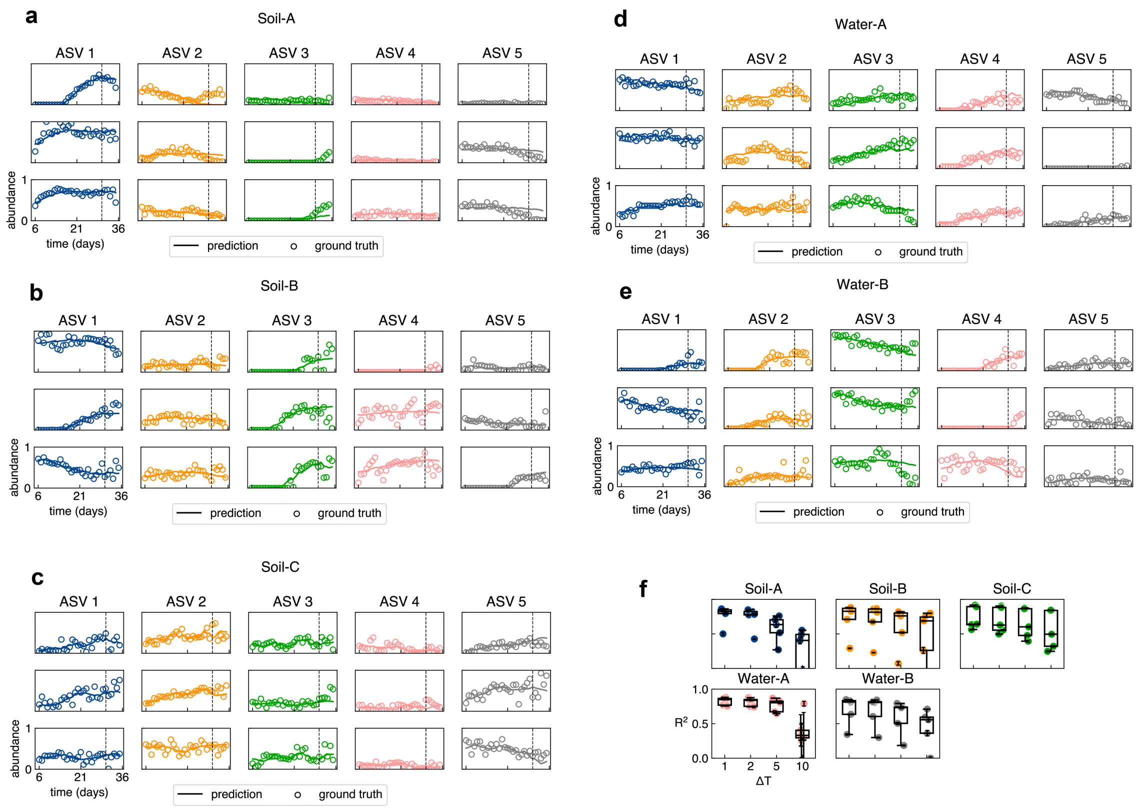


**Figure S16. Prediction of target ASV dynamics by the MLP-VAE models.**

a) – e) Example time series of predicted target ASV dynamics (curves) and ground truth (open circles) between day 6 and 35 across the five communities. Dashed line represents Day 30.

f) Forecasting accuracy ($R^2$) of ASV time series using VAE-MLP models across different time horizons. For each community (Water-A, Water-B, Soil-A, Soil-B, Soil-C), we trained models to forecast the last $\Delta T$ days of ASV trajectories ($\Delta T$ = 1, 2, 5, 10) based on the preceding 30 days. Five trials were performed for each $\Delta T$ using cross-validated MLP models in the latent space of a pretrained VAE. $R^2$ scores were computed for each of the top 5 ASVs and averaged across segments and trials.

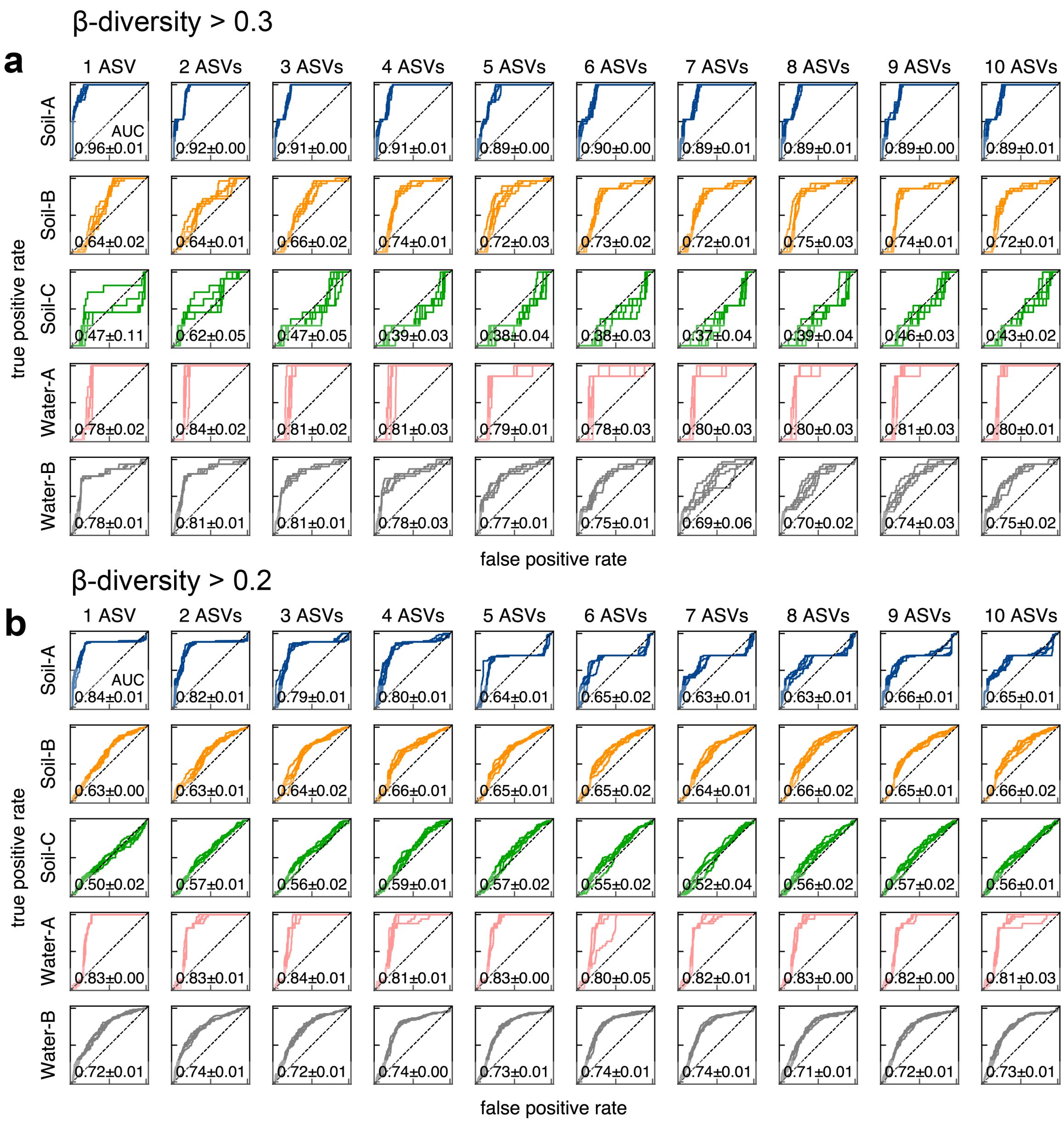


**Figure S17. Receiver Operating Characteristic (ROC) curves for classification of large community shifts.**

a) ROC curves from 5 training replicates for predicting whether β-diversity > 0.3 between day 30 and day 31, using time series data from days 1 to 30.

b) Same as (a), but for a β-diversity threshold of > 0.2.

Each panel shows performance for a specific community using a model trained on the stacked embeddings of 1 to 10 ASVs. The area under the curve (AUC, mean ± SD) is shown in each plot.

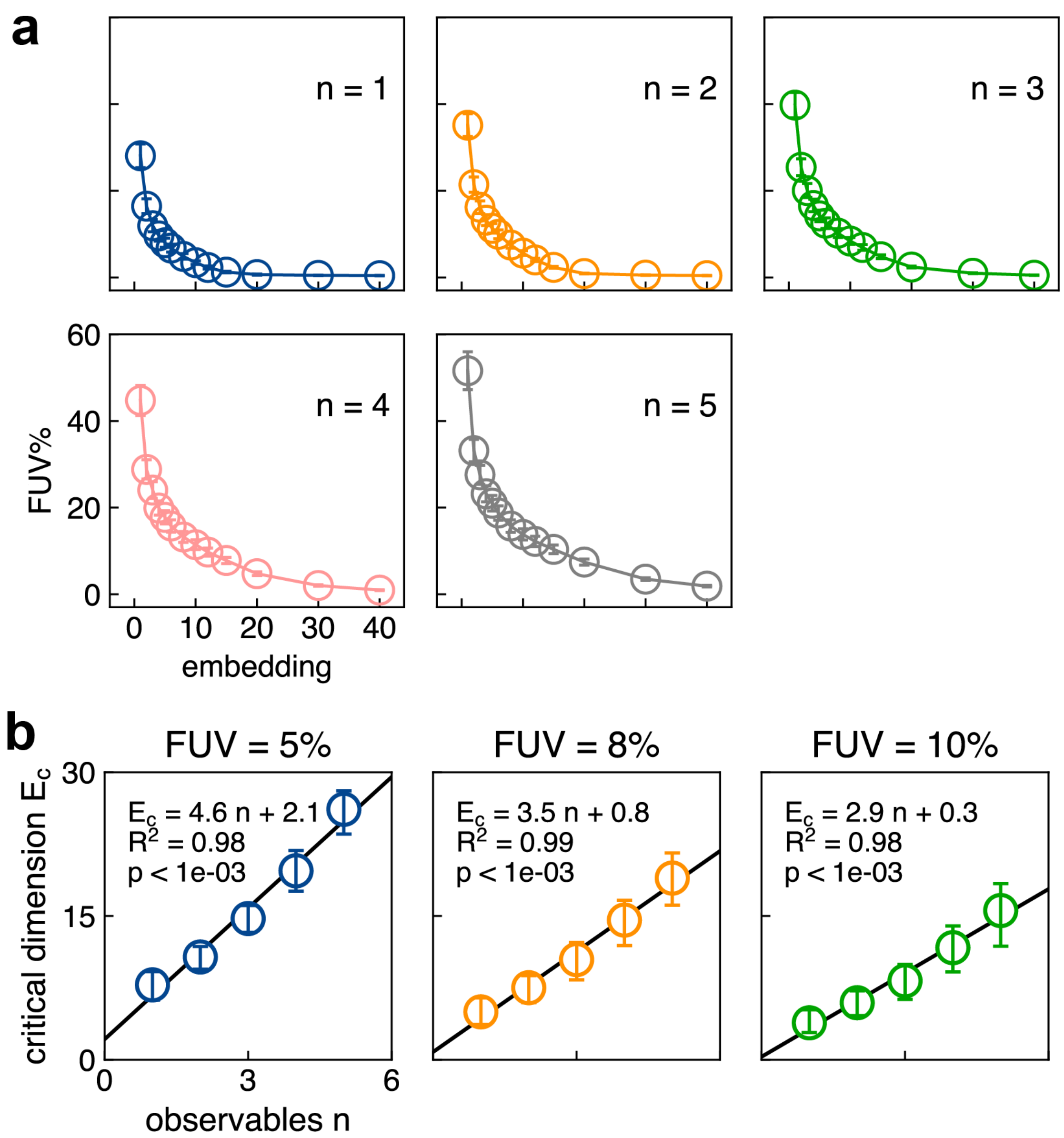


**Figure S18. Human vaginal microbiome dynamics is low dimensional.**

a) Dependence of FUV (3 training replicates are averaged for each cross-validation fold, mean ± SE across 24-fold cross-validation was shown) on the embedding dimension.

b) Linear correlations between $E_c$ and $n$ consistently holds for FUV thresholds of 5%, 8%, and 10%. Each point represents the mean $E_c$ from bootstrapping interpolation (1,000 times), together with 95% CI. Black line indicates the best linear fit to mean $E_c$

across *n*, with reported $R^2$ and bootstrapped p-value testing the slope is significantly greater than zero.

**Table S1. Keio strain library and barcodes.**

| | Barcode | Knockout gene |
|---|---|---|
| K1 (mEGFP) | ACTGACCGATATGTCAGTATGCCGTGCA | *yqeJ* |
| K2 (mTagBFP2) | CTAGACTGCAATTGCAGTACGTCGCGTA | *tag* |
| K3 (LSSmOrange) | TCGAACATGCATCTAGGTAGTCCGTCAG | *gutQ* |
| K4 (mCherry) | CTAGACAGTCATCGTAGTAGTCCGTGCA | *yfjP* |
| K5 | AGTCACATCGATCTAGGTACGTCGTGCA | *yccK* |
| K6 | AGCTACGTCAATGTCAGTTAGCCGTCGA | *yedS* |
| K7 | GACTACTACGATCTAGGTACTGCGGTCA | *sgcX* |
| K8 | TCGAACTAGCATTCGAGTTGACCGATCG | *yqiG* |
| K9 | GTCAACGACTATTCGAGTTGACCGGACT | *ybfD* |
| K10 | CTAGACGCTAATAGTCGTTCGACGCAGT | *ygcK* |
| K11 | ATCGACATCGATCTGAGTACTGCGGCAT | *narU* |
| K12 | CAGTACTCGAATTAGCGTTACGCGCATG | *ygeI* |
| K13 | TAGCACCGTAATTGCAGTTACGCGCTAG | *tolC* |
| K14 | TGCAACACGTATCTGAGTGACTCGGTAC | *yhdP* |
| K15 | TAGCACCGTAATTACGGTGATCCGGCTA | *ydcC* |
| K16 | GCTAACCGATATAGTCGTACGTCGCGTA | *yagL* |
| K17 | TCGAACGTCAATGTACGTGCATCGGATC | *yhfZ* |
| K18 | CGATACATGCATTGCAGTGACTCGAGTC | *kptA* |
| K19 | ATCGACTACGATATCGGTCTAGCGACTG | *ypjC* |
| K20 | GCATACTCAGATATGCGTGACTCGTAGC | *ygcQ* |
| K21 | ACGTACCATGATTGCAGTTACGCGGCTA | *pbl* |
| K22 | CGTAACTCAGATGATCGTATGCCGTAGC | *yhbO* |
| K23 | ATGCACGCATATGATCGTGCATCGGTAC | *yidF* |
| K24 | TCGAACTCAGATTGCAGTTACGCGATGC | *yhhI* |
| K25 | CATGACGCTAATGTACGTAGCTCGATCG | *yphB* |
| K26 | CATGACGCATATGTCAGTTGACCGCAGT | *modA* |
| K27 | ACGTACCTGAATCTAGGTTCGACGTCGA | *gudD* |
| K28 | CAGTACGTACATAGCTGTTCGACGGTAC | *phnE* |
| K29 | CTGAACAGTCATGTCAGTCAGTCGATGC | *recO* |
| K30 | CTGAACTGACATGATCGTTACGCGTCGA | *metI* |
| K31 | CGTAACACTGATTGCAGTCATGCGTCAG | *yabP* |
| K32 | CATGACCAGTATCAGTGTTAGCCGCGAT | *malT* |
| K33 | TCGAACGATCATGATCGTGATCCGGACT | *phnM* |
| K34 | CATGACAGTCATACGTGTTACGCGAGCT | *ygaT* |
| K35 | TCGAACTGACATCGATGTCTAGCGTACG | *ygcG* |
| K36 | ATCGACGCTAATAGCTGTCATGCGGCTA | *ygeN* |

| | | |
|---|---|---|
| K37 | ATGCACTACGATGACTGTAGCTCGCATG | *ybhR* |
| K38 | ACTGACCTAGATCGATGTTCGACGAGTC | *rhsC* |
| K39 | TCAGACTCGAATGATCGTCGTACGCGAT | *prpB* |
| K40 | GCATACCATGATCGTAGTCGATCGTCGA | *sseA* |
| K41 | CATGACGACTATAGCTGTATCGCGCTGA | *recN* |
| K42 | GTCAACTCAGATAGTCGTGCATCGGCAT | *ygeQ* |
| K43 | TCGAACTGCAATCTGAGTACTGCGAGCT | *yigZ* |
| K44 | TCGAACCGTAATCGATGTACGTCGCAGT | *ycdR* |
| K45 | AGTCACACTGATACGTGTGCATCGTAGC | *rhsB* |
| K46 | ACGTACCTGAATGACTGTATGCCGTACG | *ydjQ* |
| K47 | CGTAACACGTATATCGGTCTAGCGATCG | *yaiL* |
| K48 | GACTACACTGATCTGAGTCGTACGGTAC | *glyS* |
| K49 | ACGTACCATGATGATCGTAGTCCGAGCT | *ylcG* |
| K50 | AGTCACGCTAATTACGGTGATCCGTCGA | *mutS* |
| K51 | TCGAACCGATATCGTAGTGTCACGGATC | *ppdB* |
| K52 | CGATACGCTAATGCATGTGATCCGGCTA | *rpiA* |
| K53 | CTGAACGACTATATCGGTGTACCGATCG | *rhsD* |
| K54 | CATGACCATGATATGCGTCTGACGTCGA | *yeaN* |
| K55 | AGCTACTCGAATTGACGTAGCTCGTCAG | *yliH* |
| K56 | GACTACTACGATATGCGTACTGCGGCTA | *rtcB* |
| K57 | GCTAACCAGTATAGCTGTCTAGCGACGT | *rspB* |
| K58 | ATCGACGATCATCAGTGTACTGCGCATG | *dsbB* |
| K59 | TACGACACTGATGTCAGTGATCCGGCTA | *mutH* |
| K60 | GTACACTCGAATATCGGTTGCACGTGAC | *yqeH* |
| K61 | GTCAACTAGCATTCGAGTTGACCGTACG | *ygfI* |
| K62 | ACTGACTAGCATATGCGTCAGTCGCGTA | *gspD* |
